# Patterns and potential drivers of intraspecific variability in the body elemental composition of a terrestrial consumer, the snowshoe hare (*Lepus americanus*)

**DOI:** 10.1101/594101

**Authors:** Matteo Rizzuto, Shawn J. Leroux, Eric Vander Wal, Yolanda F. Wiersma, Travis R. Heckford, Juliana Balluffi-Fry

## Abstract

1. Intraspecific variability in ecological traits is widespread in nature. Recent evidence, mostly from aquatic ecosystems, shows individuals differing at the most fundamental level, that of their chemical composition. Age, sex, or body size may be key drivers of intraspecific variability in the body concentrations of carbon (C), nitrogen (N), and phosphorus (P). However, we still have a rudimentary understanding of the patterns and drivers of intraspecific variability in chemical composition of terrestrial consumers, particularly vertebrates.
2. Here, we investigate the whole-body chemical composition of snowshoe hare *Lepus americanus*, providing one of the few studies of patterns of stoichiometric variability and its potential drivers for a terrestrial vertebrate. Based on snowshoe hare ecology, we expected higher P and N concentrations in females, as well as in larger and older individuals.
3. We obtained whole-body C, N, and P concentrations and C:N, C:P, N:P ratios from a sample of 50 snowshoe hares. We then used general linear models to test for evidence of a relationship between age, sex, or body size and stoichiometric variability in hares.
4. We found considerable variation in the C, N, and P concentrations and elemental ratios within our sample. Contrary to our predictions, we found evidence of N content decreasing with age. As expected, we found evidence of P content increasing with body size. As well, we found no support for a relationship between sex and N or P content, nor for variability in C content and any of our predictor variables.
5. Despite finding considerable stoichiometric variability in our sample, we found no substantial support for age, sex, or body size to relate to this variation. The weak relationship between body N concentration and age may suggest varying nutritional requirements of individuals at different ages. Conversely, P’s weak relationship to body size appears in line with recent evidence of the potential importance of P in terrestrial systems. Snowshoe hares are a keystone herbivore in the boreal forest of North America. The substantial stoichiometric variability we find in our sample could have important implications for nutrient dynamics in both boreal and adjacent ecosystems.

## 1 Introduction

The elemental composition of an organism is an important ecological trait subject to variation within and across species (Jeyasingh *et al*., 2014; Leal *et al*., 2017). Primary producers (e.g., plants, algae), owing to the presence of dedicated storage structures in their cells, are plastic in their elemental composition (Sterner & Elser, 2002; Borer *et al*., 2013), with extreme cases in which intraspecific variation exceeds that found among species (Ågren & Weih, 2012). Marine phytoplankton and terrestrial plants show large variability in their carbon (C), nitrogen (N), and phosphorus (P) concentrations, at both large (Martiny *et al*., 2013; Sardans *et al*., 2016) and small spatiotemporal extents (Rivas-Ubach *et al*., 2012). Conversely, variability in the chemical composition of consumers is generally considered small or null, due to strict homeostasis requirements — particularly for terrestrial consumers (Sterner & Elser, 2002; Elser *et al*., 2007; Leroux & Schmitz, 2015). However, evidence for strict consumer homeostasis can be equivocal and studies of invertebrates (González *et al*., 2011) or aquatic consumers (e.g., fish; Ebel *et al*., 2015, 2016) show considerable intraspecific stoichiometric variability. For terrestrial vertebrates, much research has focused on their nutritional body composition (Hewison *et al*., 1996), differential use of chemical elements among conspecifics (Atwood & Weeks, 2002), or on their body condition (Peig & Green, 2010). We know little, however, about their organismal elemental composition, how it interacts with other ecological traits, and whether or not it varies among individuals. Knowledge of the patterns and drivers of terrestrial consumer body elemental composition may improve our ability to predict the relationship between terrestrial consumers and ecosystem function (e.g., carbon cycling).

Herbivores occupy a trophic level where they have the potential to exert top-down control on primary producers and can also affect their predators’ ecology (Leroux & Schmitz, 2015). Herbivores rely on resources whose elemental composition is markedly different from their own: plants and algae are rich in C-heavy structural molecules, while herbivores rely on N and P to fuel their growth (Fagan *et al*., 2002; Sterner & Elser, 2002). This mismatch, especially evident in terrestrial food webs, creates a strong bottleneck to nutrient flow in ecosystems (Boersma *et al*., 2008; Leroux & Schmitz, 2015). As such, investigating the drivers of intraspecific variability in elemental composition of herbivores can help shed light on both trophic dynamics and ecosystem processes, such as nutrient cycling (Sterner & Elser, 2002; Leroux & Schmitz, 2015). Previous studies showed consumers’ elemental composition varying as a function of an individual’s age, sex, or body size (Main *et al*., 1997; González *et al*., 2011; Goos *et al*., 2017). Here, we investigate how these three variables influence the whole-body elemental content of a terrestrial consumer common across North America’s boreal forest, the snowshoe hare *Lepus americanus*. We focus on C, N, and P, as these are three of the most commonly studied and important elements for an organism (Sterner & Elser, 2002, but see Jeyasingh *et al*., 2014). Owing to the strong nutrient limitation of boreal ecosystems (Pastor *et al*., 2006), and their role as keystone herbivores in them (Krebs *et al*., 2018), snowshoe hares are well-suited to address these questions.

Organismal elemental content can vary throughout an individual’s life. For instance, early life stages of *Daphnia lumholtzi* show higher concentrations of P and lower N:P than older ones, that appear to more strongly influence their growth rate than their body size (Main *et al*., 1997). Similar patterns among other phyto- and zooplankton species led to the development of the Growth Rate Hypothesis which predicts that faster growing individuals have higher body P concentrations than slower growing conspecifics, as RNA and Ribosome synthesis rely heavily on P supply (Elser *et al*., 2000; Sterner & Elser, 2002). Far from applying just to unicellular organisms, evidence shows its predictions hold true among freshwater insects as well (Back & King, 2013). Furthermore, similar intraspecific differences in elemental concentrations between life stages also exist among vertebrates (El-Sabaawi *et al*., 2012a,b, 2014). At times, this ontogenic variation in elemental composition of conspecifics is as large as that found between different genera, as is the case among minnows (*Cyprinidae* spp.; Boros *et al*., 2015). Consequently, this allows for describing life stage-specific elemental signatures, as recently done for pre- and post-spawn adult Atlantic salmon *Salmo salar* during their annual spawning migration up- and downstream, respectively (Ebel *et al*., 2016). While mammals’ life histories often do not feature dramatic events such as spawning migrations or metamorphosis, the transition from newborn to adult still involves a wide range of developmental changes, e.g., skeletal development and gonadal maturation, that could influence the elemental requirements and composition of an individual as it grows. For instance, due to its chemical composition, Sterner & Elser (2002) hypothesize that, as bone tissue should contain most of its P reserves, a vertebrate’s P content should increase with age given skeletal growth.

In a similar way, sex could affect relative content of key elements, due to the dichotomy in reproductive strategies and roles of males and females. Yet, evidence for a relationship between sex and stoichiometry is controversial. For example, costs of lactation or parental care (e.g., in bats; Hood *et al*., 2006) and development of secondary sexual characteristics (e.g., antlers; Atwood & Weeks, 2002) can influence the relative concentrations of elements (Goos *et al*., 2017). This relationship, however, is far from general. In some mayflies species, for instance, females tend to have higher %P than males and slower %P decline with age, whereas other species show no sex-related differences or even opposite trends (Back & King, 2013). Further, three-spine stickleback *Gasterosteus aculeatus* populations sampled from different lakes showed opposing trends in %P and N:P between sexes (Durston & El-Sabaawi, 2017). Finally, among guppies (*Cyprinidae* spp.), sexual differences only appeared to exert an effect on body %P when considered together with stream of origin (El-Sabaawi *et al*., 2012b). As these examples show, sex-related patterns of organismal stoichiometry and their relevance to a species’ ecology are often difficult to ascertain. As snowshoe hares are weakly sexually dimorphic (Feldhamer *et al*., 2003), and lack specialized secondary sexual characteristics, differences in the organismal content of C, N, or P could arise as a consequence of differences in body size or varying nutritional requirements between the sexes (e.g., due to gestation and lactation needs; Hood *et al*., 2006).

Organismal elemental composition can also vary with an individual’s body size, as well as with its related condition metrics (body condition indexes, BCI; Stevenson & Woods, 2006). In particular, P content tends to scale with an organism’s size (González *et al*., 2011; Back & King, 2013). While widespread, the sign of the relationship differs strongly among different groups, such as invertebrates and vertebrates. Invertebrates, lacking an internal repository of P, show a strongly negative pattern between P concentration and body size, in keeping with the GRH (Sterner & Elser, 2002; González *et al*., 2011). Conversely, as among vertebrates the majority of P stocks are found in bone tissue (Sterner & Elser, 2002), the P-body size allometric relationship should be positive. That is, P concentration should increase as individuals grow larger. However, modeling approaches show that P content should initially decreases and eventually approach an asymptotic relationship with vertebrate body size (Gillooly *et al*., 2005). Conversely, empirical evidence suggests P content increases with body size following Sterner & Elser’s prediction. Among guppies, larger individuals have higher concentrations of P than their smaller conspecifics (El-Sabaawi *et al*., 2012a). Likewise, in the Atacama Desert of Chile, two species of lizards show a similar pattern of %P increasing with body size (González *et al*., 2011). In turn, this variability in the content of fundamental nutrients with body size could influence the overall condition of an individual — which ultimately determines its fitness and nutritional value for its predators (Stevenson & Woods, 2006). In a strongly P-limited environment like the boreal forest, larger individuals could indeed show higher concentrations of P.

From all of the above it follows that, during an individual’s ontogenic development, its content of any given element of interest likely varies as a result of age (Ebel *et al*., 2016), sex (Durston & El-Sabaawi, 2017), or body size (El-Sabaawi *et al*., 2012a). Following previous works and theory (González *et al*., 2011; Boros *et al*., 2015; Ebel *et al*., 2016), we predict that (1) whole-body P content of snowshoe hares increases with increasing body size and as individuals grow older. We also expect (2) female hares to have higher content of limiting nutrients, N and P, than males, due to the higher reproductive costs. At the same time, we investigate the relationship between organismal concentration of limiting nutrients, such as N or P, and an individual’s body condition. In this case, we expect (3) snowshoe hares in better condition to have higher concentrations of N, P or both, at all life stages. We present one of the first assessments of whole-body elemental composition of a small terrestrial mammal and discuss how intraspecific stoichiometric variability might influence trophic dynamics and ecosystem processes.

## 2 Methods

### 2.1 Study Species

The snowshoe hare is the keystone herbivore in the boreal forests of North America, with a geographic range extending from Alaska to New Mexico (Feldhamer *et al*., 2003; Krebs *et al*., 2018). Average total body length of snowshoe hares varies between 36–52 cm and mean adult body weight is 1.3 kg (range: 0.9–2.3 kg), with both seasonal and annual fluctuations. Females are usually 10–25% larger than males (Feldhamer *et al*., 2003).

Snowshoe hares are mostly nocturnal and do not hibernate over winter (Feldhamer *et al*., 2003). For these reasons, they are most often found in habitats with dense understory vegetation, allowing for more effcient thermo-regulation and predator avoidance (Litvaitis *et al*., 1985). Snowshoe hares populations cycle throughout the continent, with peaks every 8–11 years and densities ranging 5 to 25 fold (Reynolds *et al*., 2017; Krebs *et al*., 2018). These abundance cycles are a defining characteristic of the boreal forest, affecting the ecology of many boreal species, from the plants the snowshoe hares consume, to their competitors and predators (Krebs *et al*., 2018).

Snowshoe hares were introduced in Newfoundland in 1864 and quickly spread across the island (Strong & Leroux, 2014). Studies conducted in the 1960s investigated their population dynamics, diet composition, and competition with another introduced herbivore, the moose *Alces alces* (Dodds, 1960, 1965). Compared to areas of Canada further west, Newfoundland has a fluctuating snowshoe hare population, with shorter and less regular periodicity (8–9 years; Reynolds *et al*., 2017). Their diet varies among seasons and areas of the island of Newfoundland (Dodds, 1960): black spruce *Picea mariana* and balsam fir *Abies balsamea* comprise most of the winter forage, whereas during the summer they feed almost exclusively on deciduous plants and shrubs (e.g., *Vaccinium* spp.; *Trifolium* spp.; *Viburnum* spp.; Dodds, 1960).

### 2.2 Data Collection

#### 2.2.1 Snowshoe hare morphology, age, and sex

In October 2016, we purchased 50 whole wild-caught snowshoe hares from a local trapper, and stored them in individual plastic bags at −20 °C. The specimens came from four trapping locations in the Eastern Avalon peninsula, over a small 21.5 km^2^ trapping area around the towns of Chapel Arm (NL, 47°31^′^00^′′^N, 53°40^′^00^′′^W) and Long Harbour (NL, 47°25^′^46^′′^ N, 53°51^′^30^′′^W). In the laboratory, we thawed and weighed each specimen to the closest 0.1 g. We collected data on total body length, left hind foot length, and skull length and width for each hare to the closest mm, repeating each measurement 3 times and using their arithmetic mean in all subsequent analyses (see Supplementary Information section S1.2).

Like rodents, the teeth of lagomorphs grow continuously during their life, making conventional aging techniques based on dentine and cement inapplicable (Morris, 1972). To account for this, we aged our specimens using a mixed approach involving counting bone tissue growth lines deposited after each winter in the mandibular bone. We used an ageing method developed for mountain hares *Lepus timidus* to select the area of the bone from which to count the growth lines (Iason, 1988). For all 50 snowshoe hares in our sample, we extracted the complete mandibular bone, cleaned it of all soft tissues, and shipped the clean bones to Matson’s Laboratory (Manhattan, MT, USA) for age determination (see SI section S1.3).

We determined specimen sex using a DNA-based approach (Shaw *et al*., 2003; see SI section S1.4). As the snowshoe hare genome is not yet completely sequenced, we used published primers for the European rabbit *Oryctolagus cuniculus* to amplify the genetic material extracted from our specimens and from two control snowshoe hares of known sex (Fontanesi *et al*., 2008). In cases when this DNA-based approach failed to detect an individual’s sex (n=3), we determined it by visual inspection and palpation of the genital area.

#### 2.2.2 Body Size Metrics

To investigate the relationship between body size and organismal chemical composition of snowshoe hares we used two different metrics: body condition and average body length. Body condition is a widely used metric to assess the overall health and quality of an animal (Stevenson & Woods, 2006; Peig & Green, 2010). To estimate body condition we used the scaled mass index (SMI; Peig & Green, 2009, 2010). The SMI standardizes an individual’s measure of body size with respect to another, thus accounting for scaling relationships (Peig & Green, 2009). In particular, the SMI uses the average value of the length measurement (*L*) with the strongest relationship with body size (i.e., its body weight, *M*) as the standardizing variable, as established by a Standardized Major Axis regression (Peig & Green, 2009; see SI section S1.5). The SMI formula is:

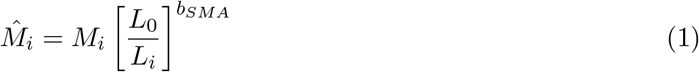

where 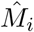 is the SMI of individual *i*, *M*_*i*_ is its body weight, *L*_*i*_ is the linear measure of body size of *i*, *b*_*SMA*_ is the exponent (i.e., slope) of a Standardized Major Axis Regression of ln(*M*) over ln(*L*), and *L*_0_ is the study population’s average value of *L*_*i*_. Therefore, the SMI is the expected weight of the individual if its length measurement was equal to the population’s average value. In this way, the SMI provides an easily understandable assessment of an animal’s condition. In this study, we used the length of the left hind foot to calculate the SMI. From the SMI value, we then computed the relative body condition (*K*_*n*_) of an individual as the ratio of *M*_*i*_ to 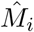 (Stevenson & Woods, 2006). This provided us with a simple metric to assess how good or bad an individual’s condition was, compared to what it should be.

As the SMI is sensitive to the length measurement used to calculate it, we ran a separate set of models using a SMI produced using skull length, which also showed a strong relationship with body weight (see SI section S1.5). Furthermore, we considered average body length as a separate estimate of the effect of body size on the C:N:P stoichiometry of snowshoe hares. We calculated average body length of individual snowshoe hares by taking the arithmetic mean of the three measurements of total body length we collected from each specimen, and used this value in all subsequent analyses.

#### 2.2.3 Whole-body Stoichiometry

After collecting both morphological data and bone samples required for ageing, we reduced the whole hare to a homogeneous paste using a Retsch GM300 knife mill (Retsch GmbH, Haan, Germany). Through preliminary tests conducted on road-killed individuals not included in our sample of 50, we noticed that elastic or fine tissues, such as skin, fur, ears, and the walls of the digestive tract, were particularly difficult to homogenize with our equipment. Consequently, we removed fur, skin, and ears from all specimens. For the digestive tract, instead, we removed, cleaned, and finely chopped it before adding it back into the mixture. For each specimen, we collected a sample of the homogenized mixture, weighed it for wet weight (g), and oven dried it for an average of 4 nights at 50 °C. After drying, we further ground each sample to as fine a powder as possible using mortar and pestle, and weighed it again for dry weight (g). On average, we required 50 g of wet homogenized material to produce 10 g of dry material for elemental composition determination. We transferred all ground samples to glass vials and stored them in desiccators to prevent moisture accumulation and mold formation.

We sent the 50 dried, whole-body samples to the Agriculture and Food Laboratory (AFL) at the University of Guelph for determination of the whole-body content of C, N, and P as % of each sample’s dry weight. At AFL, each sample was further ground before stoichiometric analyses. Concentrations of C and N were obtained by ashing the samples at 475 °C for 3 hours prior to carbon analysis using catalytic combustion (950 °C) with an Elementar vario MACRO cube (Elementar Analysensysteme GmbH, Langenselbold, Germany). This separates the desired elements from foreign gases: the elements are then analyzed using thermal conductivity detection. Organic C quantity was calculated via subtraction of inorganic C from total C obtained in this way. For P, homogenized samples were first digested with nitric acid and hydrochloric acid using a closed-vessel microwave (CEM Marsxpress, CEM Corporation, Matthews, NC, USA). The microwave-digested sample was then brought to volume with nanopure water and P content quantified using Inductively-coupled Plasma-Optical Emission Spectroscopy using a Varian Vista Pro ICP-OES and a pneumatic nebulizer (Varian Inc., Palo Alto, CA, USA) (Poitevin, 2012).

Given that few studies have homogenized and measured the elemental composition of terrestrial vertebrates, we ran pilot tests to assess within-sample variability. These showed some within-sample variability in %C and %N. To account for this, each sample was analyzed three times for C and N content. Conversely, %P was relatively invariant within samples. Because of this, only 5 samples were run in duplicate to assess within-sample variability in %P (see SI section S1.6). In addition, to capture variability within individuals due to our homogenization protocol, we selected 5 random specimens for which we sent 2 additional samples (n=10) of the homogenized paste to AFL (see SI section S1.6). Upon receiving the results back from AFL, to obtain C:N:P stoichiometry and molar ratios for each hare, we calculated each hare’s dry body weight and converted the concentration of each element to molar mass using atomic weights. As variation among samples taken from each individual was negligible for all three elements, we used average values of %C, %N and %P for each individual in subsequent analyses (see SI section S1.7).

### 2.3 Statistical Analyses

We used General Linear Models (GLMs) in R (v. 3.4.4; R Core Team, 2018) to investigate age, sex, and body size as potential drivers of whole-body hare stoichiometry. We used the concentration of each element of interest (i.e., %C, %N, %P), as well as the ratios C:N, C:P, and N:P as our response variables. Age (continuous), sex (categorical), relative body condition (*K*_*n*_, continuous), and average body length (ABL, continuous) were our explanatory variables. To test our predictions, we considered the effects of each of our predictor variables alone and their additive and 2-way interactive effects. We tested for multicollinearity among our explanatory variables using variance inflation factor analysis (VIF). As expected, VIF showed that relative body condition and average body length were highly correlated (VIF>3). Therefore we did not include these two variables in the same model (see SI section S2). We fit a set of 22 competing models, including an intercept-only model, and used the function AICc from the AICmodavg R package to select the most parsimonious model based on the Akaike Information Criterion corrected for small sample size (AICc; Burnham & Anderson, 2002; Mazerolle, 2017). We then removed models with uninformative parameters (*sensu* Arnold, 2010) from the model set of each response variable (Leroux, 2019; see SI section S3.1).

## 3 Results

Snowshoe hares in our sample varied in age between 0 (“young-of-the-year”) and 6 years old, the majority (74%) being between 0 and 1 years old. Only one individual, a female, was 6 years old. Males were more common (31 out of 50) than females (19). Average (±SD) wet body weight was 1374.81 g (±186.59, range: 914.30–1776.50 g), with average dry weight being 399.11 g (±74.70, range: 241.76–567.86 g). Water made up to 72% of body weight. Average body length was 42.49 cm (±2.07, range: 36.67–46.67 cm). Average left hind foot length (*L*_0_) for our snowshoe hare population was 12.88 cm (±0.58, range: 11.40–14.10 cm). The slope of the Standardized Major Axis Regression of average left hind foot length on body mass (i.e., the exponent *b*_*SMA*_ in Equation (1)) was 3.18. Overall, young snowshoe hares appeared more variable in relative body condition than older individuals (mean: 1.01 ±0.14; Fig. S5).

Snowshoe hares were, on average, composed of 43.60% C (±2.59, range: 37.46%–51.29%), 11.20% N (±0.78, range: 9.42%–12.68%), and 2.97% P (±0.52, range: 2.00%–4.29%; Fig. 1). The most parsimonious model for %N included only age (R^2^=0.066): %N was negatively related to the age of individual snowshoe hares (Table 1). Evidence for this relationship is, however, weak as the intercept-only model was within 2 ∆AICc of the top ranked model (Table 1). For %P, the two top ranked models included relative body condition and average body length, respectively (Table 1). %P was positively related to relative body condition (R^2^=0.073; Fig. 2) and average body length (R^2^=0.047). Again, evidence for these relationships is weak as the intercept-only model was the third-best performing model and within 2 ∆AICc of the top ranked models (Table 1). We also observed a qualitative pattern of higher %P among older males (Fig. 3), but found no statistical support for it (Table 1). For %C, the top ranked model was the intercept-only model, which provides no evidence of a relationship between variation in %C and age, sex, or body size of individuals (Table 1).

**Table 1:** Top ranking GLMs for %C, %N, and %P based on ∆AICc values. We report only models that scored better than the null model, together with the null model. k, number of parameters in the model, LL, log-likelihood, *K*_*n*_, relative body condition, ABL, average body length. We provide coeffcient values as estimate (±SE).

| %N top models |  |  |  | Coefficients |  |  |  |
| --- | --- | --- | --- | --- | --- | --- | --- |
| k | LL | $\Delta\text{AICc}$ | $R^2$ | Intercept | Age | $K_n$ | ABL |
| 3 | -56.599 | 0.000 | 0.066 | 11.367<br>($\pm 0.141$) | -0.160<br>($\pm 0.087$) | | |
| 2 | -58.306 | 1.147 | 0.000 | 11.200<br>($\pm 0.111$) | | | |
| %P top models |  |  |  | Coefficients |  |  |  |
| k | LL | $\Delta\text{AICc}$ | $R^2$ | Intercept | Age | $K_n$ | ABL |
| 3 | -35.556 | 0.000 | 0.073 | 1.962<br>($\pm 0.526$) | | 1.006<br>($\pm 0.518$) | |
| 3 | -36.252 | 1.391 | 0.047 | 0.687<br>($\pm 1.495$) | | | 0.054<br>($\pm 0.035$) |
| 2 | -37.444 | 1.508 | 0.000 | 2.974<br>($\pm 0.073$) | | | |
| %C top models |  |  |  | Coefficients |  |  |  |
| k | LL | $\Delta\text{AICc}$ | $R^2$ | Intercept | Age | $K_n$ | ABL |
| 2 | -118.090 | 0.000 | 0.000 | 43.606<br>($\pm 0.367$) | | | |

**Figure 1:**
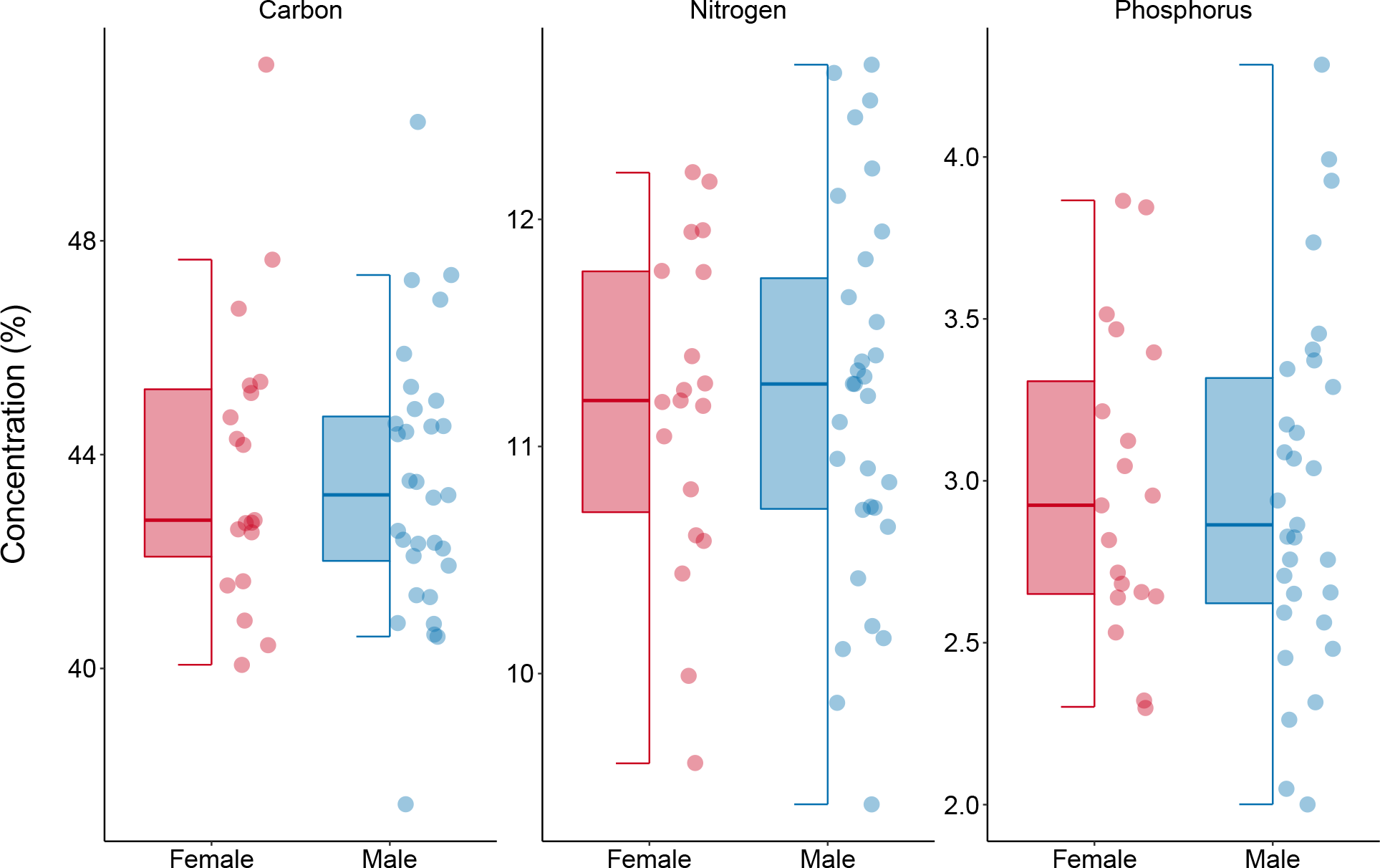
Sex-related variability in the concentrations of carbon (C), nitrogen (N), and phosphorus (P) among 50 snowshoe hares. The lower and upper boundaries of the box are the first and third quartiles, respectively. The thick horizontal line inside the box is the median, i.e., the second quartile. The whiskers extend from either boundary to no further than the largest (or smallest) value * 1.5 IQR (interquartile range). Female snowshoe hares show higher median values of %P than males. Males, on the other hand, appear consistently more variable than females in their content of both N and P. Note the different scales of the y-axis among the three panels.

**Figure 2:**
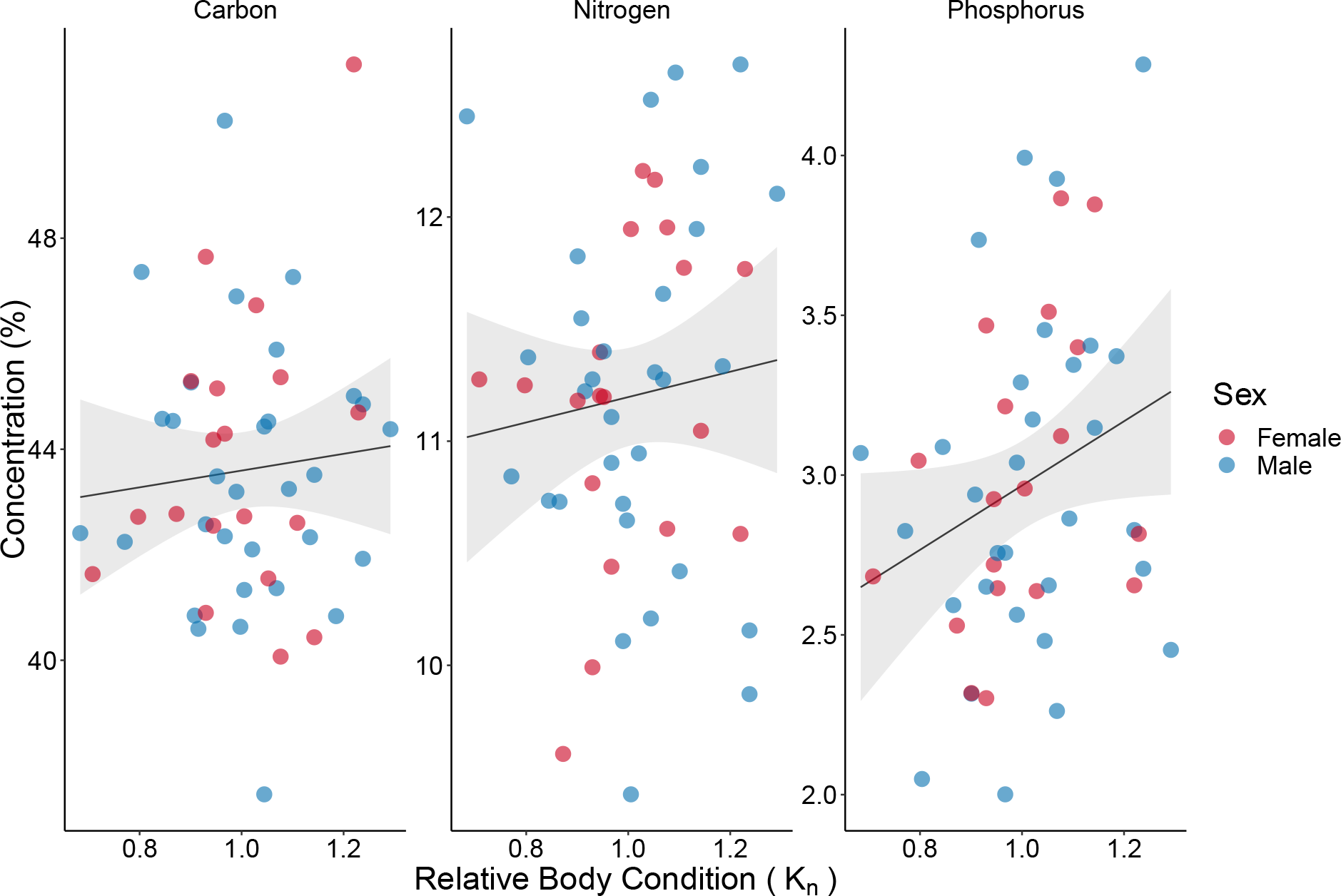
Variability in the concentrations of C, N, and P with increasing relative body condition. The positive trend for P is evident, and is weakly supported by the results of our modeling. Conversely, there is no visual evidence of a relationship between %C or %N and relative body condition, which is further confirmed by the results of our modeling (Table 1). Solid lines are ordinary least square regression lines, shaded areas represent 95% confidence intervals around them.

**Figure 3:**
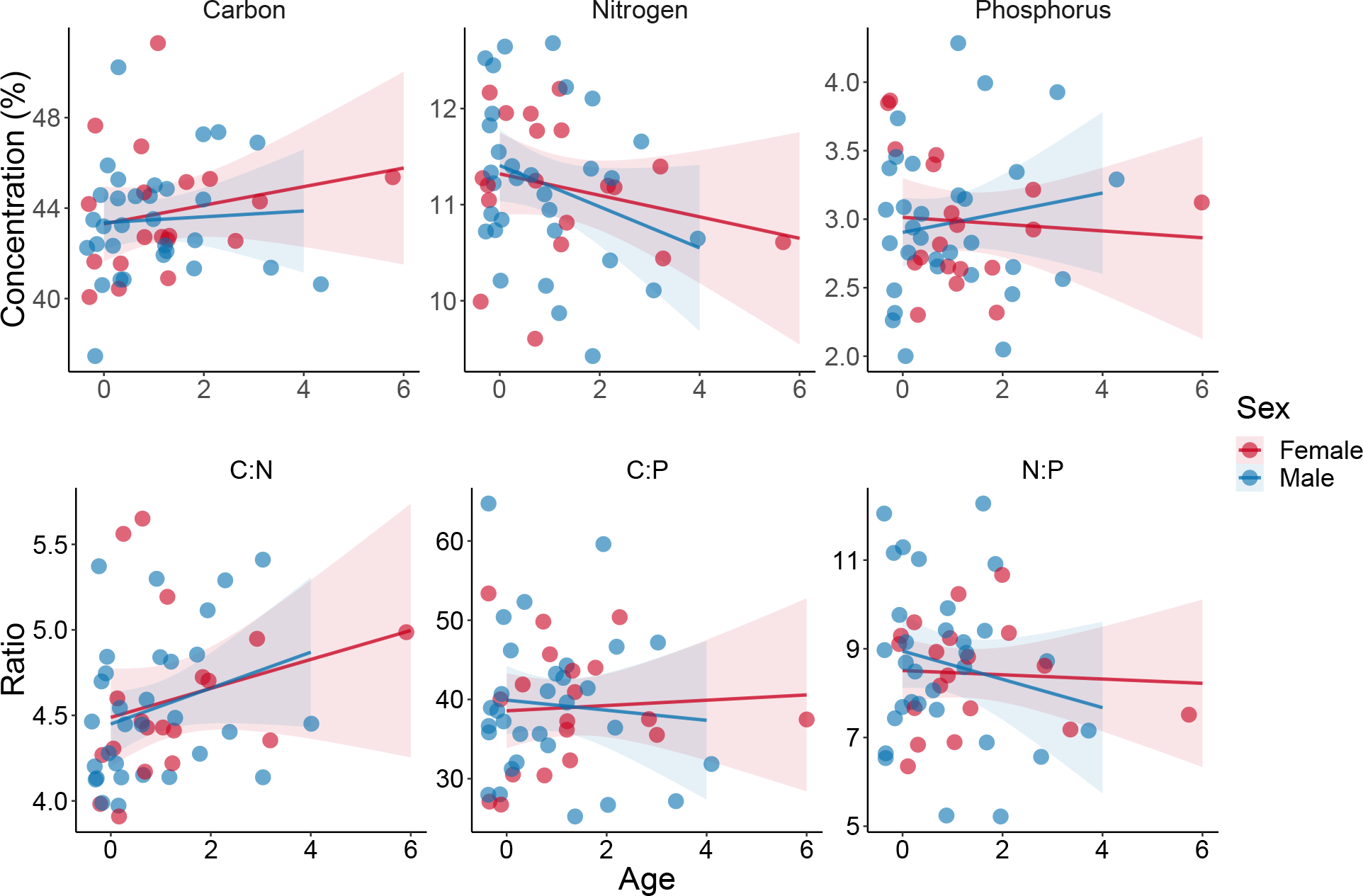
Variability in C, N, P concentrations and their stoichiometric ratios with increasing age among 50 snowshoe hares. **Upper panels**: while concentrations of P appear largely invariant as age increases, we notice a negative trend for N concentration for both sexes. This is further supported by the weak relationship found between age and %N through our modeling approach. Conversely, our modeling does not provide any support for the seemingly increasing trend we observe for %C. **Lower panels**: values of C:N appear to increase with age, for both males and females, as would be expected given the negative relationship between %N and age. Conversely, the values of N:P seem to decrease as males get older, which might mean that %N is more strongly influencing the variability of this ratio than %P is. No trend appears evident for C:P, which is in line with the lack of pattern in the variability of %C. We added a jitter to the data to improve readability of the graphs. All other specifications as in Fig. 2.

For the stoichiometric ratios, the top ranked model for C:N included only age, which had a positive relationship with C:N ratio (R^2^=0.074; Table 2). For this relationship too, evidence is weak as the intercept-only model was within 2 ∆AICc of the best-performing one. We found no evidence for a relationship between age, sex, body size, and either C:P or N:P as the top ranked model for both these ratios was the intercept-only model (Table 2). Using skull length instead of left hind foot length to calculate *K*_*n*_ did not qualitatively change our results (see SI Tables S1 and S2).

**Table 2:** Top ranking GLMs for C:N, C:P, and N:P based on ∆AICc values. All specifications as in Table 1.

| C:N top models |  |  |  | Coefficients |  |  |  |
| --- | --- | --- | --- | --- | --- | --- | --- |
| k | LL | $\Delta\text{AICc}$ | $R^2$ | Intercept | Age | $K_n$ | ABL |
| 3 | -27.818 | 0.000 | 0.074 | 4.465<br>($\pm 0.079$) | 0.095<br>($\pm 0.049$) | | |
| 2 | -29.731 | 1.559 | 0.000 | 4.564<br>($\pm 0.063$) | | | |
| C:P top models |  |  |  | Coefficients |  |  |  |
| k | LL | $\Delta\text{AICc}$ | $R^2$ | Intercept | Age | $K_n$ | ABL |
| 2 | -178.30 | 0.000 | 0.000 | 39.205<br>($\pm 1.223$) | | | |
| N:P top models |  |  |  | Coefficients |  |  |  |
| k | LL | $\Delta\text{AICc}$ | $R^2$ | Intercept | Age | $K_n$ | ABL |
| 2 | -94.153 | 0.000 | 0.000 | 8.580<br>($\pm 0.227$) | | | |

## 4 Discussion

We provide one of few assessments of the body elemental composition of a terrestrial vertebrate and investigate potential drivers of this fundamental ecological trait. Overall, we find considerable variation in the concentrations of C, N, P, and their ratios within our sample of snowshoe hares. However, age, sex, and body size appear to explain little of this variation. Our models highlight a weak and negative relationship between an individual’s age and its N concentration, and a symmetrically weak and positive trend between age and C:N. Likewise, we find weak support for a relationship between an individual’s body size and its P content. Together, these results provide some of the first evidence for intraspecific variability in the stoichiometry of a terrestrial vertebrate but raise the need to consider a broader suite of potential drivers of the variability we observed.

Based on our analyses, we found weak evidence in support of our prediction that age might drive variability in body elemental composition of snowshoe hares. In particular, we observe a negative trend in N concentration: young individuals (0–1 years old) have seemingly higher N concentrations than older ones — with a more pronounced decrease among males than among females (Fig. 3). As would be expected from this pattern, C:N values show an opposite, positive trend with age (Fig. 3) — reflecting the lower amounts of N compared to C in older hares and lending further support to this result. Age is a fundamental driver of stoichiometric differences among conspecifics, as shown for a range of different species (Boros *et al*., 2015; Ebel *et al*., 2015). Younger individuals may show higher %N as a result of increased N allocation to muscle tissue production (Boros *et al*., 2015). Snowshoe hares experience strong predation pressure from a large cohort of predators from the earliest life stages (Krebs *et al*., 2018). The higher N content among leverets we observe, then, could be a sign of early investments in production of N-rich protein to develop the muscle mass necessary for their hide and run anti-predator response. We also observed a qualitative pattern of increasing %P with age among males. While we lack quantitative support for this trend (Table 1), it is nonetheless in line with current theories. Indeed, Sterner & Elser (2002) postulate that the Growth Rate Hypothesis prediction of higher P concentrations among young, fast-growing individuals might not hold or apply differently in vertebrates, where the vast majority of P is locked in the skeleton (Sterner & Elser, 2002). Our observation of higher %P in older male hares falls in line with other recent evidence supporting Sterner & Elser’s insight: Boros *et al*. (2015), for instance, did not find an ontogenic trend in P concentration of two species of laboratory-reared minnows. Future research should assess the GRH applicability beyond the unicellular and aquatic systems in which it was originally conceived.

Counter to our prediction, we find no evidence for a relationship between hare stoichiometry and sex. Male individuals did show larger variability in their N concentration than females (Figs 1–3), but our models provide no quantitative support for this observation. This lack of evidence for differences between sexes may not be surprising. Several studies that investigated the relationship between sex and organismal stoichiometry provide contradictory evidence (El-Sabaawi *et al*., 2012b; Back & King, 2013; Goos *et al*., 2017). Among guppies, for instance, sex had no relationship with stoichiometry when considered alone, yet it had significant interactions with the fish’s stream of origin — likely an indirect consequence of different predation levels experienced by males and females in different streams (El-Sabaawi *et al*., 2012b). Conversely, a recent study on *Hyalella* amphipods found evidence of strong sexual dimorphism in the concentrations and patterns of variation of multiple elements, which underlay sexual dimorphism in traits as different as foraging behaviour, nutritional physiology, and sex-specific selection of genomic loci (Goos *et al*., 2017). Additionally, among antler-producing ungulates, males and females differ in both content and use of certain elements (e.g., calcium; Atwood & Weeks, 2002). Finally, as hares undergo morpho-physiological changes during their reproductive season, investigating the relationship between whole-body stoichiometry and sex among actively reproducing hares might produce different results (Hood *et al*., 2006). These contrasting lines of evidence highlight the need of further research, involving a wider range of species from a variety of environments, to reduce the uncertainty around the role of sex as a driver of variation in organismal stoichiometry.

Consistent with our predictions, our results indicate body size as a potential driver for variability in P concentration in our sample. The two top models for this element included relative body condition and average body length, and both variables had a positive relationship with %P. In particular, the observed body weight of snowshoe hares with higher %P matched or exceeded the predicted value obtained from the SMI formula (Equation (1)). Snowshoe hare body condition fluctuates throughout the year (Murray, 2002), with peaks in the months leading up to the boreal winter, during which hares remain active and face increased levels of stress due to both lack of optimal forage and increased predation (Krebs *et al*., 2018). As body condition declines over the winter months (Murray, 2002), it would be interesting to test whether the weak relationship we observe between P and body condition would vary in a similar way. Additionally, we observe a qualitatively larger variability in relative body condition among young hares in our sample than among older specimens (Fig. S5). Snowshoe hares produce many litters in a year (up to four; Feldhamer *et al*., 2003), yet a large number of leverets do not survive their first winter (Krebs *et al*., 2018). While we do not find evidence for a relationship between age and P content, it would nonetheless be interesting to test whether being born early or late in the year could explain part of this variability. Our results, albeit weakly supported by our statistical analyses, appear to confirm the potential role P plays within the internal chemical machinery of an animal, and its importance for its survival (Elser *et al*., 2007; Boersma *et al*., 2008).

A large amount of variability in our sample remains unexplained and, overall, we find only weak support for our initial hypothesis of ontogenic variation in organismal elemental composition among snowshoe hares. Indeed, other vertebrate species show much stronger patterns of intraspecific variation in elemental content. Ebel *et al*. (2015, 2016), for instance, showed that Atlantic salmons *S. salar* at different ontogenic stages have distinct stoichiometric signatures, particularly before and after their first migration from their freshwater nurseries to the open ocean. The reason for these differences in the magnitude of the effects mediated by ontogeny could be found in the life history of snowshoe hares. Snowshoe hares do not undergo dramatic life events like the salmon’s migration, or the metamorphosis of certain insect species, which clearly separate different life stages. Rather, they are characterized by short gestation periods (*≃*30-40 days; Feldhamer *et al*., 2003) and quick maturation of leverets into adults (*≃*6 months). It is possible, in this scenario, that we investigated the effects of age at a time in the life of snowshoe hares when most of the changes in chemical composition had already taken place. It is also interesting to note the larger proportion of young individuals in our sample, consistent with current knowledge about snowshoe hare survival beyond their first winter (Krebs *et al*., 2018). Thus, a potentially interesting and rewarding research avenue would be to further investigate differences in hare whole-body stoichiometry during the earlier stages of their lives. Finally, although our samples were collected from a small area, fine scale forage quality may also be a driver of the stoichiometric variability we observed. Future work could investigate spatial variation in habitat and forage quality as a driver of consumer body elemental composition (Leroux *et al*., 2017).

The variation in hare body composition we observe could have repercussions beyond the stoichiometry of this species, and influence ecosystem processes such as nutrient cycling, transport, and primary productivity (Pastor *et al*., 2006). Snowshoe hares are the keystone herbivore in the boreal forest, a markedly nutrient-limited environment (Pastor *et al*., 2006). They are characterized by strong, decade-long fluctuations in their population abundance and serve as primary food source for many predator species (Krebs *et al*., 2018). Paucity of nutrients, and the well-known stoichiometric mismatch between plants and herbivores (Elser *et al*., 2000; Sterner & Elser, 2002), prompted boreal forest herbivores to evolve browsing strategies allowing them to extract as much nutrients as possible from their food sources (Pastor *et al*., 2006). Thus, the appearance of a large number of young snowshoe hares over the landscape during a population peak could have strong dampening effects on elemental cycling in the boreal forest — as well as in adjacent ecosystems — possibly reducing N or P availability to primary producers as they become locked within the herbivores’ biomass. By infusing ongoing ecological research with stoichiometric data, future studies could address this potential interplay between a species’ stoichiometry and the ecosystem processes it contributes to (Leal *et al*., 2017). In turn, this would allow for shedding light on fine-grain mechanisms with far-reaching consequences, such as cross-ecosystem nutrient mobilization (Schmitz *et al*., 2018) and nutrient recycling (Schmitz *et al*., 2014), as well as on their influence on ecosystem services fundamental for humans.

Ecological stoichiometry has a long history in marine and freshwater ecosystems and has been shaped by detailed studies of algae, plants and invertebrates (Elser *et al*., 2007; González *et al*., 2011; Ågren & Weih, 2012). In recent years, researchers started investigating the stoichiometry of more complex organisms in aquatic ecosystems, particularly fish (El-Sabaawi *et al*., 2012a,b, 2014). This expanded the reach of ecological stoichiometry in exciting new directions, integrating it with other subfields of ecology, such as metabolic ecology (Rivas-Ubach *et al*., 2012), ecosystem ecology (Abbas *et al*., 2012), and landscape ecology (Sardans *et al*., 2016; Leroux *et al*., 2017). Yet, terrestrial species other than plants remain largely unexplored in terms of their stoichiometry. Our results suggest that a greater focus on terrestrial vertebrates and consumers could provide novel insights and potentially question well-known concepts in this field.

## 5 Statement of Authorship

- MR, SJL, EVW, and YFW devised the study;
- MR, TRH, JBF, SJL, YFW, and EVW collected the data;
- MR analyzed the data;
- MR, TRH, JBF, SJL, YFW, and EVW interpreted the data;
- MR led the writing of the manuscript.

All authors contributed critically to the drafts and gave final approval for publication.

## 6 Acknowledgements

The authors would like to thank Ms. Isabella Richmond for thoughtful comments on earlier drafts of the manuscript. MR would like to acknowledge the help of Mr. Shawn Reid, Ms. Meriel Fitzgerald, Mr. Benjamin Stratton, and Ms. Nathalie Djan-Chékar during data collection. MR would also like to thank Prof. Edward “Ted” Miller for pointing out the method used to age snowshoe hare. This research was funded by the Government of Newfoundland and Labrador, Innovate NL, Mitacs, the Canada Foundation for Innovation, Memorial University, and a Natural Science and Engineering Research Council Discovery Grant to SJL. This research was approved by the Memorial University Animal Care Committee, permit number 18-02-EV.

## 7 Data Availability

Data and code used in the analyses are available via the figshare online repository at: https://doi.org/10.6084/m9.figshare.7884854.v1

