## Supplementary Information for "Patterns and potential drivers of intraspecific variability in the body elemental composition of a terrestrial consumer, the snowshoe hare (*Lepus americanus*)"

Version dated: March 29, 2019

### S1 Data Collection

In this section, we provide details on the laboratory protocols we used to obtain data on age, sex, and body condition of the snowshoe hares in our sample.

#### S1.1 Data handling

We collected and handled data from our laboratory processing in digital format, by using a digital collection form (FileMaker Pro v. 14.0, FileMaker Inc., Santa Clara, CA, USA) on an iPad Mini 2 (Apple Inc., Cupertino, CA, USA), thus removing the potential error-prone step of transferring information from physical data collection forms to digital spreadsheets.

#### S1.2 Morphometric Data

Following Peig & Green (2009)'s recommendation of testing a range of different length measurements before selecting the one used in the Scaled Mass Index calculations, from each of our 50 snowshoe hares we collected four different length measurements. These were total body length, left hind foot length, skull length and skull width. We measured total body length from the tip of the nose to the anus. We measured the left hind foot from the knee joint to the tip of the nail of the middle finger, while pressing down the foot and spreading the fingers. As for the skull measurements, we took length as the distance between the tip of the nose and the base of the skull and width as the width at the cheekbones. We took each measurement to the nearest 0.1 mm and repeated each measurement three times.

#### S1.3 Age Determination

Ageing snowshoe hares can be difficult, as their teeth grow continuously throughout their lives. Hence, traditional cementum-based ageing techniques are not available for this species. We used a mixed approach that involved combining an ageing method developed by Iason (1988) for for mountain hares (*Lepus timidus*) and standard histological procedures for examining bone sections.

For each of our 50 snowshoe hares, we extracted the complete mandibular bone. We carefully cleaned it both by hand, removing as much soft tissue as possible before drying the bones out and storing them in desiccators to prevent mold formation. To further clean each bone, we let carrion beetles (family: *Dermestidae*) digest all remaining soft tissue over a period of two weeks and until the bone was completely clean. Each bone was individually tagged with its specimen's ID before beetle digestion. Once we cleaned all 50 bones, we shipped them to Matson's Laboratory (Manhattan, MT, USA) for age determination.

Here, mandible specimens were prepared for histological examination following standard procedure. Each sample was decalcified in a weak acidic solution, then embedded in a paraffin block to allow sectioning at 14 microns using a microtome. The resulting section was then mounted, stained, coverslipped, and finally examined for age determination under magnification (Figures S6 and S7).

#### S1.4 Sex Determination

Genetic sex determination followed the protocol detailed by Shaw *et al.* (2003). This protocol uses mammal-specific primers that amplify an intron region of the zinc-finger regions of the X and Y chromosomes. For snowshoe hares, these regions weight approximately 1500 bp in female (*ZFNx*) and 950 bp in males (*ZFNy*). Since no complete genome sequence exists for snowshoe hare yet, we used the set of primers for European rabbit (*Oryctolagus cuniculus*) sequenced by Fontanesi *et al.* (2008).

The PCR used a reaction volume of 25$\mu$l with 1 $\mu$l each of the forward and reverse primers and 10 $\mu$l of 2X Promega PCR Master Mix. The PCR consisted of one 5-minutes cycle at 95 °C. This was followed by a sequence of 30 s at 94 °C, 60 s at 52 °C, and 60 s at 72 °C, which was repeated for 35 cycles before a final cycle at 72 °C for 2 minutes. The end product was held at 4 °C before being electrophoresed on a 1.5% agarose gel in 1X TBE. The Genomics and Proteomics Laboratory at Memorial University of Newfoundland performed all DNA- based analyses and determined the sex of the individuals.

#### S1.5 Scaled Mass Index Calculation

Multiple indices of body condition exist which differ in both how they define "body condition" and in how they calculate it (Peig & Green, 2010). Most indices fall in one of three categories: ratios, whose units are often difficult to interpret; residuals, computed using units of mass; or non-dimensional indices (Stevenson & Woods, 2006). Recent evidence also suggests that multiple regression, while not an index *per se*, provides a valuable alternative (Labocha *et al.*, 2014). As the debate surrounding BCIs is ongoing, no clear "best" index emerged so far, and the decision on which one to apply to one's work is often heavily dependent on the history and traditions of a certain subfield of ecology (Stevenson & Woods, 2006; Peig & Green, 2010). IN our study, we used the Scaled Mass Index developed by Peig & Green (2009).

We computed the Scaled Mass Index (SMI) following the procedure detailed by Peig & Green (2009), which consists of three steps: (1) investigate which body length measurement ($L$) has the strongest relationship with body weight ($M$) using bivariate plots, (2) fit a Standardized Major Axis (SMA) regression to the ln-transformed bivariate plots, and (3) calculate the SMI using the scaling exponent of the SMA ($b_{SMA}$) from the strongest length-weight relationship identified earlier and a Thorpe-Lleonart scaling model (see main text for details). We began by producing bivariate plots for each of the four length measurements we collected from our specimens (Figure S1). From visual inspection, both average left hind foot length (HFL) and average skull length appeared to have a strong relationship with both body weight. HFL also correlated strongly with average body length (ABL), as well as allowing us to compare our results with other published studies. Thus, we chose to proceed using HFL. We then used the function `sma` in R package `smatr` (Warton *et al.*, 2012) to fit a SMA to the ln-transformed values of HFL and body weight. Finally, we extracted the slope value ($b_{SMA}$) and computed the SMI using Peig & Green's equation (Equation (1)).

As the SMI presents a number of individual components that vary, we tested the sensitivity of our results to changes in the length measurement used to calculate it by running the analyses again using Skull Length instead of HFL. We obtained qualitatively similar results with this alternative measure of body size (Tables S1 and S2).

#### S1.6 Intra-individual Stoichiometric Variability

Our study is one of the first to specifically assess intraspecific variability in the content of C, N, and P in a terrestrial vertebrate. As such, no precedent existed that could inform us as to whether our hares could show significant intra-individual variability in the concentrations of the three elements of interest or not. We addressed this issue in two ways.

First, during our laboratory sample collection, we randomly selected five individuals. For each of these specimens, we collected three separate samples of the homogeneous paste resulting from

**Table S1:** Top ranking GLMs for %C, %N, and %P based on ΔAICc, when Scale Mass Index and relative body condition ($K_n$) calculations are based on average skull length. Only the models that scored better than the null model are reported, together with the null model. k, number of parameters in a model, LL, log-likelihood, $SkK_n$, skull length-derived relative body condition, ABL, average total body length. Coefficient values are presented as estimate (±SE).

| **%N top models** | | | | **Coefficients** | | | |
|---|---|---|---|---|---|---|---|
| k | LL | ΔAICc | $R^2$ | Intercept | Age | $SkK_n$ | ABL |
| 3 | -56.599 | 0.000 | 0.066 | 11.367 (±0.141) | -0.160 (±0.087) | | |
| 2 | -58.306 | 1.147 | 0.000 | 11.200 (±0.111) | | | |

| **%P top models** | | | | **Coefficients** | | | |
|---|---|---|---|---|---|---|---|
| k | LL | ΔAICc | $R^2$ | Intercept | Age | $SkK_n$ | ABL |
| 3 | -36.25192 | 0.000 | 0.047 | 0.687 (±1.495) | | | 0.054 (±0.035) |
| 2 | -37.444 | 0.118 | 0.000 | 2.974 (±0.073) | | | |

| **%C top models** | | | | **Coefficients** | | | |
|---|---|---|---|---|---|---|---|
| k | LL | ΔAICc | $R^2$ | Intercept | Age | $SkK_n$ | ABL |
| 2 | -118.090 | 0.000 | 0.000 | 43.606 (±0.367) | | | |

**Table S2:** Top ranking GLMs for C:N, C:P, and N:P values based on ΔAICc, when Scale Mass Index and relative body condition ($K_n$) calculations are based on average skull length. All specification as in Table S1.

| **C:N top models** | | | | **Coefficients** | | | |
|---|---|---|---|---|---|---|---|
| k | LL | ΔAICc | $R^2$ | Intercept | Age | $SkK_n$ | ABL |
| 3 | -27.818 | 0.00 | 0.074 | 4.465 (±0.079) | 0.095 (±0.049) | | |
| 2 | -29.731 | 1.59 | 0.000 | 4.564 (±0.063) | | | |
| **C:P top models** | | | | **Coefficients** | | | |
| k | LL | ΔAICc | $R^2$ | Intercept | Age | $SkK_n$ | ABL |
| 2 | -178.30 | 0.000 | 0.000 | 39.205 (±1.223) | | | |
| **N:P top models** | | | | **Coefficients** | | | |
| k | LL | ΔAICc | $R^2$ | Intercept | Age | $SkK_n$ | ABL |
| 2 | -94.153 | 0.000 | 0.000 | 8.580 (±0.227) | | | |

our homogenization process. These three samples were identified with a progressive letter appended to their individual identifier (e.g., "TCH037_A", "TCH037_B", "TCH037_C") and underwent the same drying, hand-grinding, weighting, storing, and analysis protocol as the rest of the samples. Thus, along the 50 samples we sent to the Agriculture and Food Laboratory at the University of Guelph, we sent 10 additional samples, 2 each for specimens TCH037, TCH040, TCH042, TCH045, and TCH048. Second, at AFL, lab technicians ran the analyses in triplicate on each sample, providing us with a quantitative assessment of within-sample variability. We adopted this approach because pilot analyses performed at AFL on a subset of samples indicated that %C and %N had a potentially higher intra-individual variability than %P. For %P, we nonetheless ran 5 samples in duplicate as part of AFL's internal quality assurance protocol. The raw data we received from AFL are available online as a separate dataset and shown in Figures S2 and S3. None of our samples presented strong intra-individual variability in the content of C and N, so we averaged the three %C and %N values we received from each samples in subsequent analyses. As for the samples we submitted to AFL in triplicate, we computed the grand mean (mean of the mean) of the values of %C and %N, and then used these values in the analyses.

##### S1.7 Obtaining Molar Weights and Elemental Ratios

To obtain molar and stoichiometric ratios for the three elements of interest, we converted the original percentage data into weight of each element. To do this, we first calculated the dry body weight of each snowshoe hare in our sample as:

$$\frac{Sample\ Dry\ Weight}{Sample\ Wet\ Weight} : \frac{Hare\ Dry\ Weight}{Hare\ Wet\ Weight} \tag{S1}$$

We then used the atomic weights for Carbon (C), Nitrogen (N), and Phosphorus (P) to calculate the corresponding molar ratios (Meija *et al.*, 2016). We then computed the stoichiometric ratios by dividing the molar ratio of two elements, and repeated this procedure for each pair of elements. As part of this process, we also calculated each hare's water content (in g) as:

$$\left(\frac{Sample\ Wet\ Weight - Sample\ Dry\ Weight}{Sample\ Wet\ Weight}\right) \times Hare\ Wet\ Weight \tag{S2}$$

#### S2 Variance Inflation Factor Analysis

We used Variance Inflation Factor analysis to investigate collinearity among the predictor variables included in our models. We tested for independence among our variables using a VIF threshold value of <3 (Yalcin & Leroux, 2018). To do so, we used function `vifstep` in R package `usdm` (Naimi *et al.*, 2014), and ran the analyses twice, once for each of the two length measurements used to calculated the SMI and $K_n$ (see above, Section S1.5). The results of the VIF analyses indicate that, when using HFL to calculated the SMI, average body length has a collinearity problem (i.e., VIF>3). We addressed this problem by never fitting a model containing $K_n$ and ABL at the same time. In all other cases, no collinearity issues arise (Table S3).

#### S3 Model Selection

##### S3.1 Removal of Uninformative Parameters

An uninformative parameter (or "pretending variable") is a variable that does not have a relationship with the response, does not improve a model's fit to the data (i.e., its log-likelihood) or does so

**Table S3:** Results of the Variance Inflation Factor analysis run on the four explanatory variables included in the set of 22 GLMs fitted to the data. Note that no model included both Relative Body Condition ($K_n$) and Average Body Length (ABL). The $> 3$ result for ABL when SMI, and hence $K_n$, is calculated from HFL is likely due to the stronger relationship between ABL and HFL than between ABL and Skull Length.

| **Variable** | **Did it pass the VIF <3 test?** | |
|---|---|---|
| | SMI from HFL | SMI from Skull Length |
| $K_n$ | 2.504 | 2.101 |
| Age | 1.572 | 1.432 |
| Sex | 1.034 | 1.064 |
| ABL | **3.225** | 2.052 |

only marginally but, based on its AICc value, is included in a model that is ranked close to models with informative parameters (Burnham & Anderson, 2002; Arnold, 2010; Leroux, 2019). Reporting and interpretation of results from models including uninformative parameters is a widespread, yet unappreciated, issue in ecological literature (Leroux, 2019). To avoid this issue, after fitting our set of models to each response variable, we reviewed the resulting AICc table and removed models including likely uninformative parameters. We followed Leroux (2019)'s decision tree to identify and deal with uninformative parameters in our model set. We report a summarized version of each response's AICc table in the main text. Below, we report the complete AICc tables for each response variable, after removal of models containing uninformative parameters.

Tables S4 to S9 show the modeling results when using left hind foot length to calculate the relative body condition ($K_n$). For %N, $K_n$, sex, and average body condition (ABL) were uninformative (Table S4). For %P, both sex and age behaved as pretending variables and we thus removed them from the final AICc table (Table S5). For %C, all parameters were uninformative (Table S6). For the C:N ratio, all variables other than age behaved as uninformative parameters (Table S7). For C:P and N:P ratios, we found that all variables were uninformative parameters (Tables S8 and S9).

Tables S10 to S15 show the results from models including relative body condition calculated using skull length as length measurement in the SMI formula (Sk$K_n$; see Equation (1) in main text). In this scenario, for %N, all variables other than age proved to be uninformative parameters and we removed all models including them (Table S10). For %P, we found that the age, sex, and relative body condition were uninformative parameters, and thus we removed them (Table S11). For %C, we found all parameters to be pretending variables (Table S12). For the C:N ratio, all variables other than age were uninformative parameters (Table S13). For C:P and N:P ratios, we removed all models other than the null model, as all variables were uninformative (Tables S14 and S15).

**Table S4:** Full AICc table for the model set fitted to the %N data. Only one model performed better than the intercept-only (i.e., null) model. The table is sorted according to the smallest ΔAICc value. In this case, relative body condition ($K_n$) was calculated using left hind foot length (see main text). For each model, we report the number of parameters it estimates (k), its AICc and ΔAICc values, the model's Log-Likelihood (LL) and the model's fit to the data ($R^2$). Relative body condition ($K_n$), sex, and average body length (ABL) were uninformative parameters. Accordingly, we greyed-out all models including these variables.

| Model | k | AICc | ΔAICc | LL | $R^2$ |
|---|---|---|---|---|---|
| Age | 3 | 119.721 | 0.000 | -56.600 | 0.066 |
| Intercept | 2 | 120.868 | 1.147 | -58.306 | 0.000 |
| Age + $K_n$ | 4 | 121.194 | 1.473 | -56.153 | 0.083 |
| Age + ABL | 4 | 121.735 | 2.014 | -56.423 | 0.073 |
| Age + Sex | 4 | 122.076 | 2.355 | -56.594 | 0.066 |
| $K_n$ | 3 | 122.633 | 2.912 | -58.055 | 0.010 |
| ABL | 3 | 122.985 | 3.264 | -58.232 | 0.003 |
| Sex | 3 | 123.112 | 3.391 | -58.295 | 0.000 |
| Age + $K_n$ + Age:$K_n$ | 5 | 123.473 | 3.753 | -56.055 | 0.086 |
| Age + $K_n$ + Sex | 5 | 123.632 | 3.911 | -56.134 | 0.083 |
| Age + ABL + Age:ABL | 5 | 124.016 | 4.295 | -56.326 | 0.076 |
| Age + Sex + Age:Sex | 5 | 124.193 | 4.472 | -56.415 | 0.073 |
| Age + ABL + Sex | 5 | 124.196 | 4.475 | -56.416 | 0.073 |
| Sex + $K_n$ | 4 | 124.989 | 5.269 | -58.050 | 0.010 |
| Sex + ABL | 4 | 125.335 | 5.614 | -58.223 | 0.003 |
| Age + $K_n$ + Sex + Sex:$K_n$ | 6 | 125.762 | 6.041 | -55.904 | 0.092 |
| Age + $K_n$ + Sex + Age:$K_n$ | 6 | 126.006 | 6.285 | -56.026 | 0.087 |
| Age + ABL + Sex + Sex:ABL | 6 | 126.423 | 6.703 | -56.235 | 0.080 |
| Age + ABL + Sex + Age:ABL | 6 | 126.594 | 6.873 | -56.320 | 0.076 |
| Sex + $K_n$ + Sex:$K_n$ | 5 | 126.965 | 7.244 | -57.801 | 0.020 |
| Sex + ABL + Sex:ABL | 5 | 127.730 | 8.009 | -58.183 | 0.005 |
| Age + $K_n$ + Sex + Sex:$K_n$ + Age:$K_n$ | 7 | 128.261 | 8.540 | -55.797 | 0.096 |
| Age + ABL + Sex + Sex:ABL + Age:ABL | 7 | 129.038 | 9.317 | -56.186 | 0.081 |

**Table S5:** Full AICc table for the model set fitted to the %P data. Two models performed better than the intercept-only (i.e., null) model. Both include proxies for body size: relative body condition and average body length, respectively. Both sex and age were uninformative parameters. All specifications as in Table S4.

| Model | k | AICc | ΔAICc | LL | $R^2$ |
|---|---|---|---|---|---|
| $K_n$ | 3 | 77.635 | 0.000 | -35.556 | 0.073 |
| ABL | 3 | 79.026 | 1.391 | -36.252 | 0.047 |
| Intercept | 2 | 79.144 | 1.509 | -37.444 | 0.000 |
| Sex + $K_n$ | 4 | 79.946 | 2.312 | -35.529 | 0.074 |
| Age + $K_n$ | 4 | 79.970 | 2.335 | -35.540 | 0.073 |
| Age + ABL | 4 | 81.208 | 3.573 | -36.159 | 0.050 |
| Age | 3 | 81.266 | 3.632 | -37.372 | 0.003 |
| Sex + ABL | 4 | 81.393 | 3.758 | -36.252 | 0.047 |
| Sex | 3 | 81.402 | 3.767 | -37.440 | 0.000 |
| Sex + $K_n$ + Sex:$K_n$ | 5 | 82.328 | 4.693 | -35.482 | 0.075 |
| Age + $K_n$ + Sex | 5 | 82.400 | 4.765 | -35.518 | 0.074 |
| Age + $K_n$ + Age:$K_n$ | 5 | 82.403 | 4.768 | -35.519 | 0.074 |
| Age + ABL + Age:ABL | 5 | 82.905 | 5.270 | -35.771 | 0.065 |
| Sex + ABL + Sex:ABL | 5 | 83.583 | 5.949 | -36.110 | 0.052 |
| Age + Sex | 4 | 83.632 | 5.997 | -37.372 | 0.003 |
| Age + ABL + Sex | 5 | 83.678 | 6.043 | -36.157 | 0.050 |
| Age + $K_n$ + Sex + Sex:$K_n$ | 6 | 84.894 | 7.259 | -35.470 | 0.076 |
| Age + $K_n$ + Sex + Age:$K_n$ | 6 | 84.938 | 7.303 | -35.492 | 0.075 |
| Age + Sex + Age:Sex | 5 | 85.418 | 7.783 | -37.027 | 0.017 |
| Age + ABL + Sex + Age:ABL | 6 | 85.492 | 7.858 | -35.769 | 0.065 |
| Age + ABL + Sex + Sex:ABL | 6 | 86.045 | 8.410 | -36.046 | 0.054 |
| Age + $K_n$ + Sex + Sex:$K_n$ + Age:$K_n$ | 7 | 87.556 | 9.921 | -35.445 | 0.077 |
| Age + ABL + Sex + Sex:ABL + Age:ABL | 7 | 87.733 | 10.098 | -35.533 | 0.074 |

**Table S6:** Full AICc table for the model set fitted to the %C data. No model performed better than the intercept-only (i.e., null) model. All parameters were uninformative. All other specifications as in Table S4.

| Model | k | AICc | ΔAICc | LL | $R^2$ |
|---|---|---|---|---|---|
| Intercept | 2 | 240.436 | 0.000 | -118.090 | 0.000 |
| Age | 3 | 241.703 | 1.268 | -117.591 | 0.020 |
| $K_n$ | 3 | 242.340 | 1.905 | -117.909 | 0.007 |
| ABL | 3 | 242.452 | 2.016 | -117.965 | 0.005 |
| Sex | 3 | 242.484 | 2.048 | -117.981 | 0.004 |
| Age + $K_n$ | 4 | 243.820 | 3.384 | -117.465 | 0.025 |
| Age + Sex | 4 | 243.959 | 3.523 | -117.535 | 0.022 |
| Age + ABL | 4 | 244.070 | 3.635 | -117.591 | 0.020 |
| Sex + $K_n$ | 4 | 244.445 | 4.009 | -117.778 | 0.012 |
| Sex + ABL | 4 | 244.624 | 4.188 | -117.868 | 0.009 |
| Sex + $K_n$ + Sex:$K_n$ | 5 | 245.787 | 5.351 | -117.211 | 0.035 |
| Sex + ABL + Sex:ABL | 5 | 245.986 | 5.551 | -117.311 | 0.031 |
| Age + $K_n$ + Sex | 5 | 246.150 | 5.714 | -117.393 | 0.027 |
| Age + $K_n$ + Age:$K_n$ | 5 | 246.163 | 5.727 | -117.399 | 0.027 |
| Age + Sex + Age:Sex | 5 | 246.197 | 5.761 | -117.416 | 0.027 |
| Age + ABL + Sex | 5 | 246.433 | 5.997 | -117.535 | 0.022 |
| Age + ABL + Age:ABL | 5 | 246.473 | 6.038 | -117.555 | 0.021 |
| Age + $K_n$ + Sex + Sex:$K_n$ | 6 | 247.534 | 7.098 | -116.790 | 0.051 |
| Age + ABL + Sex + Sex:ABL | 6 | 248.131 | 7.695 | -117.089 | 0.039 |
| Age + $K_n$ + Sex + Age:$K_n$ | 6 | 248.576 | 8.140 | -117.311 | 0.031 |
| Age + ABL + Sex + Age:ABL | 6 | 248.947 | 8.511 | -117.497 | 0.023 |
| Age + $K_n$ + Sex + Sex:$K_n$ + Age:$K_n$ | 7 | 250.083 | 9.647 | -116.708 | 0.054 |
| Age + ABL + Sex + Sex:ABL + Age:ABL | 7 | 250.595 | 10.159 | -116.964 | 0.044 |

**Table S7:** Full AICc table for the model set fitted to the C:N data. The age-only model was the only one that performed better than the intercept-only (i.e., null) model. All other parameters were uninformative. All specifications as in Table S4.

| Model | k | AICc | ΔAICc | LL | $R^2$ |
|---|---|---|---|---|---|
| Age | 3 | 62.158 | 0.000 | -27.818 | 0.074 |
| Intercept | 2 | 63.718 | 1.559 | -29.731 | 0.000 |
| Age + ABL | 4 | 64.391 | 2.233 | -27.751 | 0.076 |
| Age + $K_n$ | 4 | 64.452 | 2.293 | -27.781 | 0.075 |
| Age + Sex | 4 | 64.507 | 2.349 | -27.809 | 0.074 |
| ABL | 3 | 65.587 | 3.428 | -29.533 | 0.008 |
| Sex | 3 | 65.825 | 3.667 | -29.652 | 0.003 |
| $K_n$ | 3 | 65.982 | 3.823 | -29.730 | 0.000 |
| Age + ABL + Age:ABL | 5 | 66.668 | 4.510 | -27.652 | 0.080 |
| Age + ABL + Sex | 5 | 66.849 | 4.690 | -27.743 | 0.076 |
| Age + $K_n$ + Sex | 5 | 66.914 | 4.755 | -27.775 | 0.075 |
| Age + $K_n$ + Age:$K_n$ | 5 | 66.926 | 4.768 | -27.781 | 0.075 |
| Age + Sex + Age:Sex | 5 | 66.938 | 4.779 | -27.787 | 0.075 |
| Sex + ABL | 4 | 67.820 | 5.661 | -29.465 | 0.011 |
| Sex + $K_n$ | 4 | 68.192 | 6.033 | -29.652 | 0.003 |
| Age + ABL + Sex + Age:ABL | 6 | 69.238 | 7.080 | -27.642 | 0.080 |
| Age + ABL + Sex + Sex:ABL | 6 | 69.419 | 7.261 | -27.733 | 0.077 |
| Age + $K_n$ + Sex + Sex:$K_n$ | 6 | 69.478 | 7.320 | -27.762 | 0.076 |
| Age + $K_n$ + Sex + Age:$K_n$ | 6 | 69.503 | 7.345 | -27.775 | 0.075 |
| Sex + ABL + Sex:ABL | 5 | 70.121 | 7.963 | -29.379 | 0.014 |
| Sex + $K_n$ + Sex:$K_n$ | 5 | 70.657 | 8.498 | -29.646 | 0.003 |
| Age + ABL + Sex + Sex:ABL + Age:ABL | 7 | 71.892 | 9.733 | -27.613 | 0.081 |
| Age + $K_n$ + Sex + Sex:$K_n$ + Age:$K_n$ | 7 | 72.191 | 10.033 | -27.762 | 0.076 |

**Table S8:** Full AICc table for the model set fitted to the C:P data. No model performed better than the intercept-only (i.e., null) model. All parameters were uninformative. All specifications as in Table S4.

| Model | k | AICc | ΔAICc | LL | $R^2$ |
|---|---|---|---|---|---|
| Intercept | 2 | 360.864 | 0.000 | -178.304 | 0.000 |
| $K_n$ | 3 | 361.583 | 0.719 | -177.531 | 0.030 |
| ABL | 3 | 361.821 | 0.957 | -177.650 | 0.026 |
| Age | 3 | 363.107 | 2.243 | -178.293 | 0.000 |
| Sex | 3 | 363.111 | 2.247 | -178.294 | 0.000 |
| Sex + $K_n$ | 4 | 363.897 | 3.033 | -177.504 | 0.032 |
| Age + $K_n$ | 4 | 363.950 | 3.086 | -177.531 | 0.030 |
| Age + ABL | 4 | 363.970 | 3.106 | -177.541 | 0.030 |
| Sex + ABL | 4 | 364.182 | 3.318 | -177.646 | 0.026 |
| Age + Sex | 4 | 365.460 | 4.596 | -178.285 | 0.001 |
| Age + ABL + Age:ABL | 5 | 365.689 | 4.825 | -177.163 | 0.045 |
| Sex + ABL + Sex:ABL | 5 | 366.249 | 5.385 | -177.443 | 0.034 |
| Age + $K_n$ + Age:$K_n$ | 5 | 366.294 | 5.430 | -177.465 | 0.033 |
| Sex + $K_n$ + Sex:$K_n$ | 5 | 366.351 | 5.487 | -177.494 | 0.032 |
| Age + $K_n$ + Sex | 5 | 366.371 | 5.507 | -177.504 | 0.032 |
| Age + ABL + Sex | 5 | 366.425 | 5.561 | -177.531 | 0.030 |
| Age + Sex + Age:Sex | 5 | 367.688 | 6.824 | -178.162 | 0.006 |
| Age + ABL + Sex + Age:ABL | 6 | 368.263 | 7.399 | -177.155 | 0.045 |
| Age + ABL + Sex + Sex:ABL | 6 | 368.688 | 7.824 | -177.367 | 0.037 |
| Age + $K_n$ + Sex + Age:$K_n$ | 6 | 368.811 | 7.947 | -177.429 | 0.034 |
| Age + $K_n$ + Sex + Sex:$K_n$ | 6 | 368.940 | 8.077 | -177.494 | 0.032 |
| Age + ABL + Sex + Sex:ABL + Age:ABL | 7 | 370.355 | 9.492 | -176.844 | 0.057 |
| Age + $K_n$ + Sex + Sex:$K_n$ + Age:$K_n$ | 7 | 371.503 | 10.639 | -177.418 | 0.035 |

**Table S9:** Full AICc table for the model set fitted to the N:P data. No model performed better than the intercept-only (i.e., null) model. All parameters were uninformative. All specifications as in Table S4.

| Model | k | AICc | ΔAICc | LL | $R^2$ |
|---|---|---|---|---|---|
| Intercept | 2 | 192.561 | 0.000 | -94.153 | 0.000 |
| ABL | 3 | 192.627 | 0.066 | -93.053 | 0.043 |
| $K_n$ | 3 | 193.015 | 0.454 | -93.247 | 0.036 |
| Age | 3 | 193.747 | 1.186 | -93.613 | 0.021 |
| Age + $K_n$ | 4 | 194.546 | 1.985 | -92.829 | 0.052 |
| Sex | 3 | 194.618 | 2.056 | -94.048 | 0.004 |
| Sex + ABL | 4 | 194.844 | 2.282 | -92.978 | 0.046 |
| Age + ABL | 4 | 194.856 | 2.294 | -92.983 | 0.046 |
| Sex + $K_n$ | 4 | 195.064 | 2.502 | -93.088 | 0.042 |
| Age + Sex | 4 | 196.012 | 3.450 | -93.561 | 0.023 |
| Age + ABL + Age:ABL | 5 | 196.760 | 4.199 | -92.698 | 0.057 |
| Age + $K_n$ + Sex | 5 | 196.830 | 4.269 | -92.733 | 0.055 |
| Age + $K_n$ + Age:$K_n$ | 5 | 196.953 | 4.392 | -92.795 | 0.053 |
| Sex + ABL + Sex:ABL | 5 | 197.122 | 4.561 | -92.879 | 0.050 |
| Age + ABL + Sex | 5 | 197.213 | 4.652 | -92.925 | 0.048 |
| Sex + $K_n$ + Sex:$K_n$ | 5 | 197.539 | 4.977 | -93.087 | 0.042 |
| Age + Sex + Age:Sex | 5 | 197.915 | 5.354 | -93.276 | 0.034 |
| Age + ABL + Sex + Age:ABL | 6 | 199.242 | 6.681 | -92.644 | 0.059 |
| Age + $K_n$ + Sex + Age:$K_n$ | 6 | 199.326 | 6.764 | -92.686 | 0.057 |
| Age + $K_n$ + Sex + Sex:$K_n$ | 6 | 199.419 | 6.857 | -92.733 | 0.055 |
| Age + ABL + Sex + Sex:ABL | 6 | 199.552 | 6.991 | -92.799 | 0.053 |
| Age + ABL + Sex + Sex:ABL + Age:ABL | 7 | 201.485 | 8.924 | -92.409 | 0.067 |
| Age + $K_n$ + Sex + Sex:$K_n$ + Age:$K_n$ | 7 | 202.038 | 9.476 | -92.686 | 0.057 |

**Table S10:** Full AICc table for the model set fitted to the %N data. In this case, relative body condition was calculated using skull length (Sk$K_n$; see text for details). Only one model performed better than the intercept-only (i.e., null) model. Sex, average body length, and skull length-derived relative body condition were uninformative parameters. All other specifications as in Table S4.

| Model | k | AICc | ΔAICc | LL | $R^2$ |
|---|---|---|---|---|---|
| Age | 3 | 119.721 | 0.000 | -56.600 | 0.066 |
| Intercept | 2 | 120.868 | 1.147 | -58.306 | 0.000 |
| Age + ABL | 4 | 121.735 | 2.014 | -56.423 | 0.073 |
| Age + Sk$K_n$ | 4 | 121.981 | 2.260 | -56.546 | 0.068 |
| Age + Sex | 4 | 122.076 | 2.355 | -56.594 | 0.066 |
| Sk$K_n$ | 3 | 122.773 | 3.052 | -58.126 | 0.007 |
| ABL | 3 | 122.985 | 3.264 | -58.232 | 0.003 |
| Sex | 3 | 123.112 | 3.391 | -58.295 | 0.000 |
| Age + Sk$K_n$ + Age:Sk$K_n$ | 5 | 123.845 | 4.124 | -56.241 | 0.079 |
| Age + ABL + Age:ABL | 5 | 124.016 | 4.295 | -56.326 | 0.076 |
| Age + Sex + Age:Sex | 5 | 124.193 | 4.472 | -56.415 | 0.073 |
| Age + ABL + Sex | 5 | 124.196 | 4.475 | -56.416 | 0.073 |
| Age + Sk$K_n$ + Sex | 5 | 124.427 | 4.707 | -56.532 | 0.069 |
| Sex + Sk$K_n$ | 4 | 125.099 | 5.378 | -58.105 | 0.008 |
| Sex + ABL | 4 | 125.335 | 5.614 | -58.223 | 0.003 |
| Age + Sk$K_n$ + Sex + Sex:Sk$K_n$ | 6 | 125.756 | 6.035 | -55.901 | 0.092 |
| Sex + Sk$K_n$ + Sex:Sk$K_n$ | 5 | 126.389 | 6.668 | -57.513 | 0.031 |
| Age + ABL + Sex + Sex:ABL | 6 | 126.423 | 6.703 | -56.235 | 0.080 |
| Age + Sk$K_n$ + Sex + Age:Sk$K_n$ | 6 | 126.432 | 6.711 | -56.239 | 0.079 |
| Age + ABL + Sex + Age:ABL | 6 | 126.594 | 6.873 | -56.320 | 0.076 |
| Sex + ABL + Sex:ABL | 5 | 127.730 | 8.009 | -58.183 | 0.005 |
| Age + Sk$K_n$ + Sex + Sex:Sk$K_n$ + Age:Sk$K_n$ | 7 | 128.018 | 8.298 | -55.676 | 0.100 |
| Age + ABL + Sex + Sex:ABL + Age:ABL | 7 | 129.038 | 9.317 | -56.186 | 0.081 |

**Table S11:** Full AICc table for the model set fitted to the %P data. In this case, relative body condition was calculated using skull length (Sk$K_n$; see text for details). Only one model performed better than the intercept-only (i.e., null) model. Age, sex, and skull length-derived relative body condition were uninformative parameters. All other specifications as in Table S4.

| Model | k | AICc | ΔAICc | LL | $R^2$ |
|---|---|---|---|---|---|
| ABL | 3 | 79.026 | 0.000 | -36.252 | 0.047 |
| Intercept | 2 | 79.144 | 0.118 | -37.444 | 0.000 |
| Age + ABL | 4 | 81.208 | 2.182 | -36.159 | 0.050 |
| Age | 3 | 81.266 | 2.241 | -37.372 | 0.003 |
| Sk$K_n$ | 3 | 81.364 | 2.339 | -37.421 | 0.001 |
| Sex + ABL | 4 | 81.393 | 2.367 | -36.252 | 0.047 |
| Sex | 3 | 81.402 | 2.376 | -37.440 | 0.000 |
| Age + ABL + Age:ABL | 5 | 82.905 | 3.879 | -35.771 | 0.065 |
| Sex + ABL + Sex:ABL | 5 | 83.583 | 4.558 | -36.110 | 0.052 |
| Age + Sex | 4 | 83.632 | 4.606 | -37.372 | 0.003 |
| Age + Sk$K_n$ | 4 | 83.632 | 4.607 | -37.372 | 0.003 |
| Age + ABL + Sex | 5 | 83.678 | 4.652 | -36.157 | 0.050 |
| Sex + Sk$K_n$ | 4 | 83.719 | 4.694 | -37.415 | 0.001 |
| Age + Sex + Age:Sex | 5 | 85.418 | 6.392 | -37.027 | 0.017 |
| Age + ABL + Sex + Age:ABL | 6 | 85.492 | 6.467 | -35.769 | 0.065 |
| Age + Sk$K_n$ + Age:Sk$K_n$ | 5 | 85.822 | 6.797 | -37.229 | 0.009 |
| Age + ABL + Sex + Sex:ABL | 6 | 86.045 | 7.020 | -36.046 | 0.054 |
| Age + Sk$K_n$ + Sex | 5 | 86.105 | 7.079 | -37.371 | 0.003 |
| Sex + Sk$K_n$ + Sex:Sk$K_n$ | 5 | 86.189 | 7.164 | -37.413 | 0.001 |
| Age + ABL + Sex + Sex:ABL + Age:ABL | 7 | 87.733 | 8.707 | -35.533 | 0.074 |
| Age + Sk$K_n$ + Sex + Age:Sk$K_n$ | 6 | 88.370 | 9.344 | -37.208 | 0.009 |
| Age + Sk$K_n$ + Sex + Sex:Sk$K_n$ | 6 | 88.690 | 9.664 | -37.368 | 0.003 |
| Age + Sk$K_n$ + Sex + Sex:Sk$K_n$ + Age:Sk$K_n$ | 7 | 91.083 | 12.057 | -37.208 | 0.009 |

**Table S12:** Full AICc table for the model set fitted to the %C data. In this case, relative body condition was calculated using skull length (Sk$K_n$; see text for details). No model performed better than the intercept-only (i.e., null) model, and all parameters were uninformative. All other specifications as in Table S4.

| Model | k | AICc | ΔAICc | LL | $R^2$ |
|---|---|---|---|---|---|
| Intercept | 2 | 240.436 | 0.000 | -118.090 | 0.000 |
| Sk$K_n$ | 3 | 241.628 | 1.192 | -117.553 | 0.021 |
| Age | 3 | 241.703 | 1.268 | -117.591 | 0.020 |
| ABL | 3 | 242.452 | 2.016 | -117.965 | 0.005 |
| Sex | 3 | 242.484 | 2.048 | -117.981 | 0.004 |
| Age + Sk$K_n$ | 4 | 243.665 | 3.229 | -117.388 | 0.028 |
| Sex + Sk$K_n$ | 4 | 243.680 | 3.244 | -117.396 | 0.027 |
| Age + Sex | 4 | 243.959 | 3.523 | -117.535 | 0.022 |
| Age + ABL | 4 | 244.070 | 3.635 | -117.591 | 0.020 |
| Sex + ABL | 4 | 244.624 | 4.188 | -117.868 | 0.009 |
| Sex + Sk$K_n$ + Sex:Sk$K_n$ | 5 | 245.690 | 5.254 | -117.163 | 0.036 |
| Age + Sk$K_n$ + Age:Sk$K_n$ | 5 | 245.772 | 5.336 | -117.204 | 0.035 |
| Age + Sk$K_n$ + Sex | 5 | 245.934 | 5.498 | -117.285 | 0.032 |
| Sex + ABL + Sex:ABL | 5 | 245.986 | 5.551 | -117.311 | 0.031 |
| Age + Sex + Age:Sex | 5 | 246.197 | 5.761 | -117.416 | 0.027 |
| Age + ABL + Sex | 5 | 246.433 | 5.997 | -117.535 | 0.022 |
| Age + ABL + Age:ABL | 5 | 246.473 | 6.038 | -117.555 | 0.021 |
| Age + Sk$K_n$ + Sex + Sex:Sk$K_n$ | 6 | 248.057 | 7.621 | -117.052 | 0.041 |
| Age + ABL + Sex + Sex:ABL | 6 | 248.131 | 7.695 | -117.089 | 0.039 |
| Age + Sk$K_n$ + Sex + Age:Sk$K_n$ | 6 | 248.275 | 7.839 | -117.161 | 0.037 |
| Age + ABL + Sex + Age:ABL | 6 | 248.947 | 8.511 | -117.497 | 0.023 |
| Age + Sk$K_n$ + Sex + Sex:Sk$K_n$ + Age:Sk$K_n$ | 7 | 250.577 | 10.141 | -116.955 | 0.044 |
| Age + ABL + Sex + Sex:ABL + Age:ABL | 7 | 250.595 | 10.159 | -116.964 | 0.044 |

**Table S13:** Full AICc table for the model set fitted to the C:N data. In this case, relative body condition was calculated using skull length (Sk$K_n$; see text for details). Only one model performed better than the intercept-only (i.e., null) model. Sex, average body length, and skull length-derived relative body condition were uninformative parameters. All other specifications as in Table S4.

| Model | k | AICc | ΔAICc | LL | $R^2$ |
|---|---|---|---|---|---|
| Age | 3 | 62.158 | 0.000 | -27.818 | 0.074 |
| Intercept | 2 | 63.718 | 1.559 | -29.731 | 0.000 |
| Age + ABL | 4 | 64.391 | 2.233 | -27.751 | 0.076 |
| Age + Sk$K_n$ | 4 | 64.456 | 2.297 | -27.783 | 0.075 |
| Age + Sex | 4 | 64.507 | 2.349 | -27.809 | 0.074 |
| Sk$K_n$ | 3 | 64.646 | 2.487 | -29.062 | 0.026 |
| ABL | 3 | 65.587 | 3.428 | -29.533 | 0.008 |
| Sex | 3 | 65.825 | 3.667 | -29.652 | 0.003 |
| Age + ABL + Age:ABL | 5 | 66.668 | 4.510 | -27.652 | 0.080 |
| Sex + Sk$K_n$ | 4 | 66.760 | 4.601 | -28.935 | 0.031 |
| Age + ABL + Sex | 5 | 66.849 | 4.690 | -27.743 | 0.076 |
| Age + Sk$K_n$ + Age:Sk$K_n$ | 5 | 66.863 | 4.704 | -27.750 | 0.076 |
| Age + Sk$K_n$ + Sex | 5 | 66.896 | 4.738 | -27.766 | 0.076 |
| Age + Sex + Age:Sex | 5 | 66.938 | 4.779 | -27.787 | 0.075 |
| Sex + ABL | 4 | 67.820 | 5.661 | -29.465 | 0.011 |
| Sex + Sk$K_n$ + Sex:Sk$K_n$ | 5 | 69.015 | 6.856 | -28.826 | 0.036 |
| Age + ABL + Sex + Age:ABL | 6 | 69.238 | 7.080 | -27.642 | 0.080 |
| Age + Sk$K_n$ + Sex + Sex:Sk$K_n$ | 6 | 69.257 | 7.098 | -27.652 | 0.080 |
| Age + Sk$K_n$ + Sex + Age:Sk$K_n$ | 6 | 69.380 | 7.221 | -27.713 | 0.078 |
| Age + ABL + Sex + Sex:ABL | 6 | 69.419 | 7.261 | -27.733 | 0.077 |
| Sex + ABL + Sex:ABL | 5 | 70.121 | 7.963 | -29.379 | 0.014 |
| Age + Sk$K_n$ + Sex + Sex:Sk$K_n$ + Age:Sk$K_n$ | 7 | 71.889 | 9.731 | -27.611 | 0.081 |
| Age + ABL + Sex + Sex:ABL + Age:ABL | 7 | 71.892 | 9.733 | -27.613 | 0.081 |

**Table S14:** Full AICc table for the model set fitted to the C:P data, when using skull length to calculate relative body condition (Sk$K_n$). No model performed better than the intercept-only (i.e., null) model. All parameters were uninformative. All specifications as in Table S4.

| Model | k | AICc | ΔAICc | LL | $R^2$ |
|---|---|---|---|---|---|
| Intercept | 2 | 360.864 | 0.000 | -178.304 | 0.000 |
| ABL | 3 | 361.821 | 0.957 | -177.650 | 0.026 |
| Age | 3 | 363.107 | 2.243 | -178.293 | 0.000 |
| Sex | 3 | 363.111 | 2.247 | -178.294 | 0.000 |
| Sk$K_n$ | 3 | 363.124 | 2.260 | -178.301 | 0.000 |
| Age + ABL | 4 | 363.970 | 3.106 | -177.541 | 0.030 |
| Sex + ABL | 4 | 364.182 | 3.318 | -177.646 | 0.026 |
| Age + Sex | 4 | 365.460 | 4.596 | -178.285 | 0.001 |
| Sex + Sk$K_n$ | 4 | 365.469 | 4.605 | -178.290 | 0.001 |
| Age + Sk$K_n$ | 4 | 365.474 | 4.610 | -178.293 | 0.000 |
| Age + ABL + Age:ABL | 5 | 365.689 | 4.825 | -177.163 | 0.045 |
| Sex + ABL + Sex:ABL | 5 | 366.249 | 5.385 | -177.443 | 0.034 |
| Age + ABL + Sex | 5 | 366.425 | 5.561 | -177.531 | 0.030 |
| Age + Sk$K_n$ + Age:Sk$K_n$ | 5 | 367.687 | 6.823 | -178.162 | 0.006 |
| Age + Sex + Age:Sex | 5 | 367.688 | 6.824 | -178.162 | 0.006 |
| Sex + Sk$K_n$ + Sex:Sk$K_n$ | 5 | 367.925 | 7.061 | -178.281 | 0.001 |
| Age + Sk$K_n$ + Sex | 5 | 367.933 | 7.070 | -178.285 | 0.001 |
| Age + ABL + Sex + Age:ABL | 6 | 368.263 | 7.399 | -177.155 | 0.045 |
| Age + ABL + Sex + Sex:ABL | 6 | 368.688 | 7.824 | -177.367 | 0.037 |
| Age + Sk$K_n$ + Sex + Age:Sk$K_n$ | 6 | 370.198 | 9.335 | -178.122 | 0.007 |
| Age + ABL + Sex + Sex:ABL + Age:ABL | 7 | 370.355 | 9.492 | -176.844 | 0.057 |
| Age + Sk$K_n$ + Sex + Sex:Sk$K_n$ | 6 | 370.504 | 9.640 | -178.275 | 0.001 |
| Age + Sk$K_n$ + Sex + Sex:Sk$K_n$ + Age:Sk$K_n$ | 7 | 372.904 | 12.040 | -178.119 | 0.007 |

**Table S15:** Full AICc table for the model set fitted to the N:P data, when using skull length to calculate relative body condition (Sk$K_n$). No model performed better than the intercept-only (i.e., null) model. All parameters were uninformative. All specifications as in Table S4.

| Model | k | AICc | ΔAICc | LL | $R^2$ |
|---|---|---|---|---|---|
| Intercept | 2 | 192.561 | 0.000 | -94.153 | 0.000 |
| ABL | 3 | 192.627 | 0.066 | -93.053 | 0.043 |
| Age | 3 | 193.747 | 1.186 | -93.613 | 0.021 |
| Sk$K_n$ | 3 | 194.596 | 2.034 | -94.037 | 0.005 |
| Sex | 3 | 194.618 | 2.056 | -94.048 | 0.004 |
| Sex + ABL | 4 | 194.844 | 2.282 | -92.978 | 0.046 |
| Age + ABL | 4 | 194.856 | 2.294 | -92.983 | 0.046 |
| Age + Sex | 4 | 196.012 | 3.450 | -93.561 | 0.023 |
| Age + Sk$K_n$ | 4 | 196.114 | 3.552 | -93.612 | 0.021 |
| Sex + Sk$K_n$ | 4 | 196.711 | 4.150 | -93.911 | 0.010 |
| Age + ABL + Age:ABL | 5 | 196.760 | 4.199 | -92.698 | 0.057 |
| Sex + ABL + Sex:ABL | 5 | 197.122 | 4.561 | -92.879 | 0.050 |
| Age + ABL + Sex | 5 | 197.213 | 4.652 | -92.925 | 0.048 |
| Age + Sk$K_n$ + Age:Sk$K_n$ | 5 | 197.867 | 5.306 | -93.252 | 0.035 |
| Age + Sex + Age:Sex | 5 | 197.915 | 5.354 | -93.276 | 0.034 |
| Age + Sk$K_n$ + Sex | 5 | 198.485 | 5.924 | -93.561 | 0.023 |
| Sex + Sk$K_n$ + Sex:Sk$K_n$ | 5 | 199.060 | 6.499 | -93.848 | 0.012 |
| Age + ABL + Sex + Age:ABL | 6 | 199.242 | 6.681 | -92.644 | 0.059 |
| Age + ABL + Sex + Sex:ABL | 6 | 199.552 | 6.991 | -92.799 | 0.053 |
| Age + Sk$K_n$ + Sex + Age:Sk$K_n$ | 6 | 200.111 | 7.549 | -93.079 | 0.042 |
| Age + Sk$K_n$ + Sex + Sex:Sk$K_n$ | 6 | 200.948 | 8.387 | -93.497 | 0.026 |
| Age + ABL + Sex + Sex:ABL + Age:ABL | 7 | 201.485 | 8.924 | -92.409 | 0.067 |
| Age + Sk$K_n$ + Sex + Sex:Sk$K_n$ + Age:Sk$K_n$ | 7 | 202.752 | 10.190 | -93.042 | 0.043 |

### S4 Additional Figures

In this section, we provide additional figures and graphs. Figure S1 shows the bivariate plots used to select the best length measurement to calculate the SMI. Figures S2 and S3 show the amount of intra-individual variability in the C and N content of hares in our sample. Figure S4 shows the variability in P concentration found among three repeated samples taken from 5 random snowshoe hares. Figure S5 shows variability in relative body condition ($K_n$) and average body length among different hares of different age and sex. Figures S6 and S7 provide examples of the mandibular bone sections used to age snowshoe hares in our sample. Figure S6 shows the section obtained by the oldest individual in our sample, a 6 years old male, whereas Figure S7 show the same section but for a 1 year old hare.

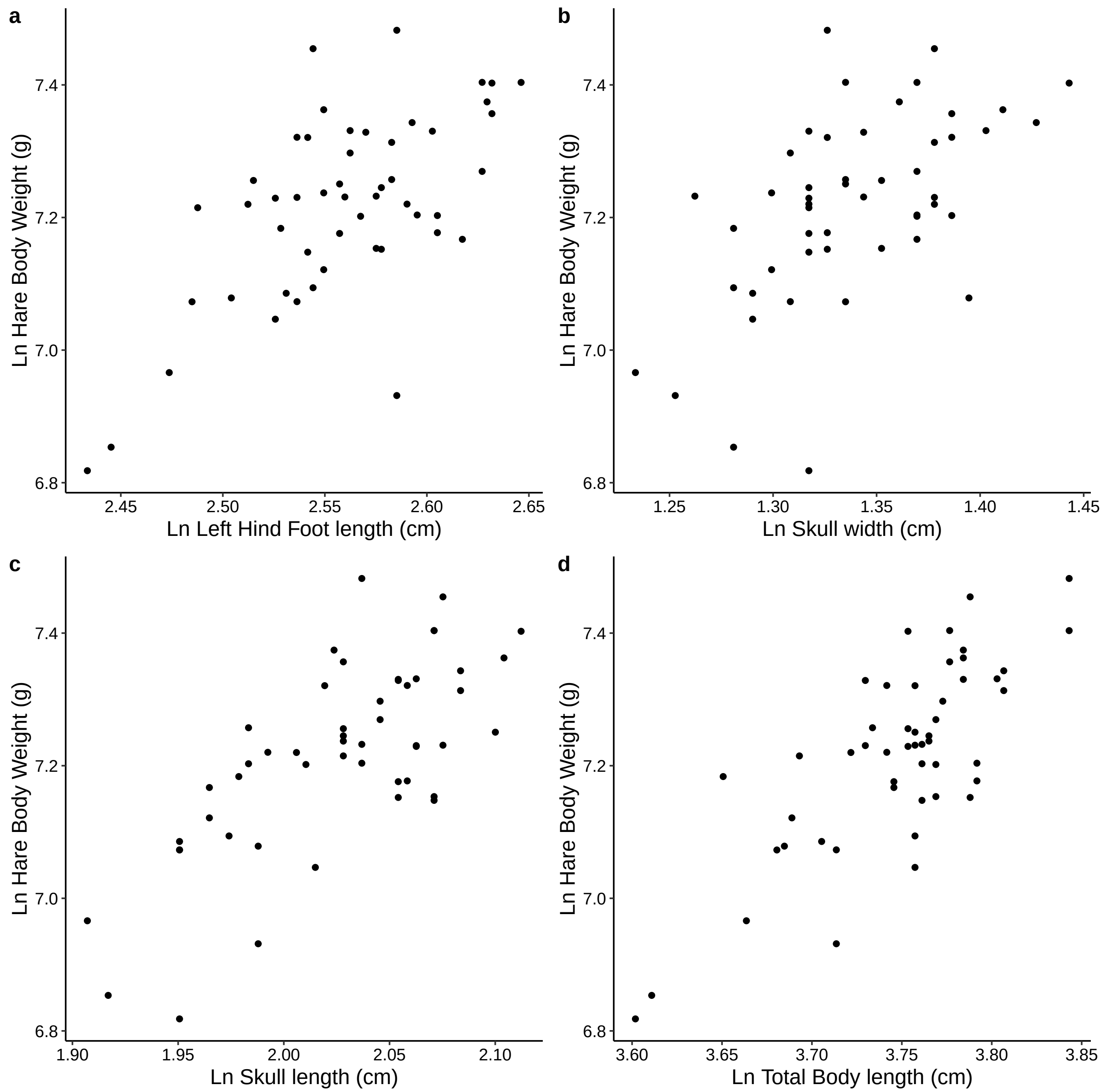

**Figure S1:** Bivariate plots used to select the length measurement to calculate the SMI. On the y-axis is the ln-transformed hare body weight in g. The x- axis reports each ln-transformed length measurement: (**a**) left hind foot length, (**b**) skull width, (**c**) skull length, and (**d**) total body length. All length measurements are in mm. Note the different scales of each x-axis. Each data point is the arithmetic mean of three measurements repeated on a single specimen.

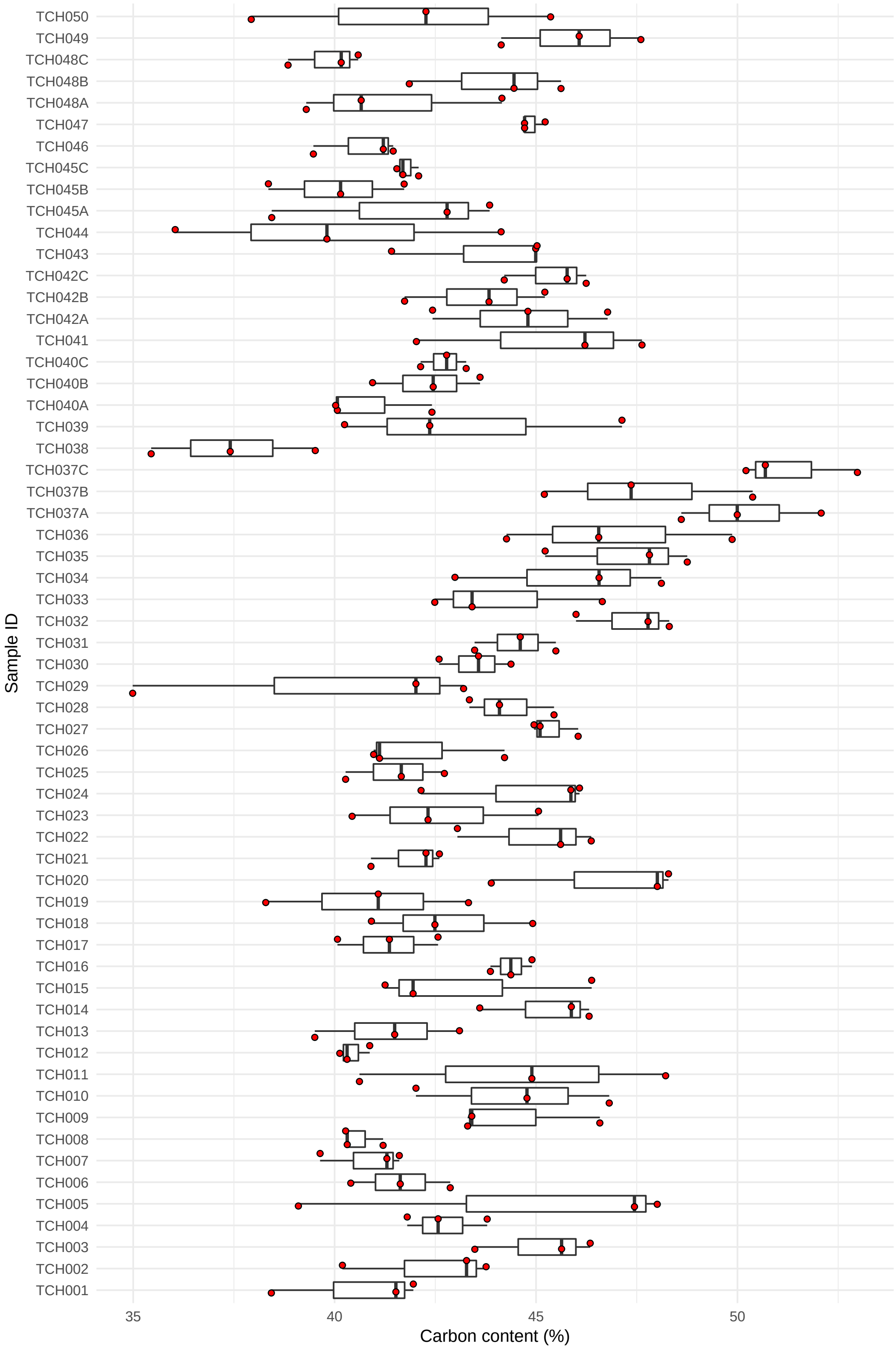

**Figure S2:** Within-sample variability in the concentration of C among the 50 snowshoe hare samples sent to AFL. The dark line inside the box is the median, the upper and lower hinges represent the 75th and 25th percentile respectively, and the two whiskers extend to 1.5 times the distance between the first and third quartile.

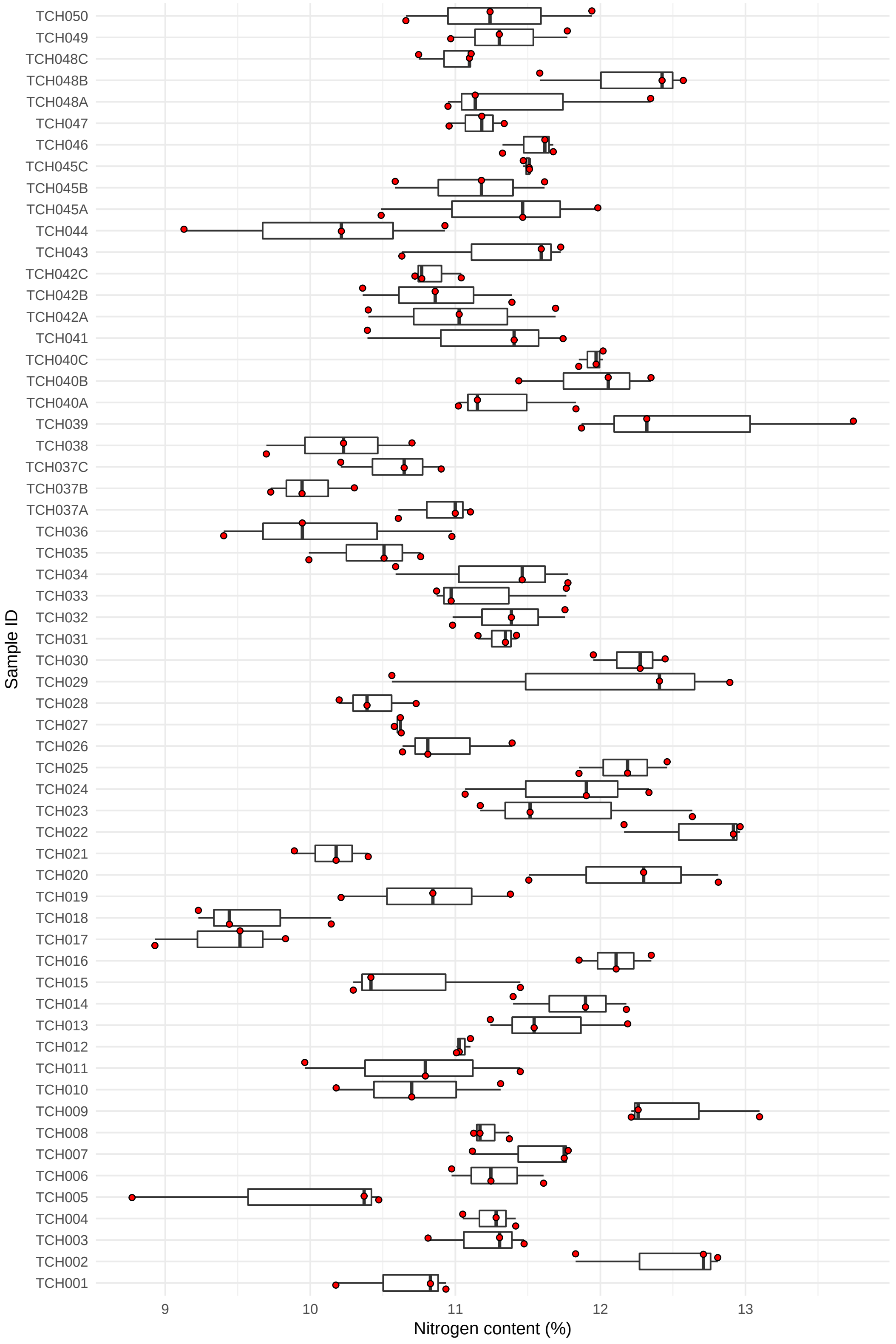

**Figure S3:** Within-sample variability in the concentration of N among the 50 snowshoe hare samples sent to AFL. All specifications as in Figure S2.

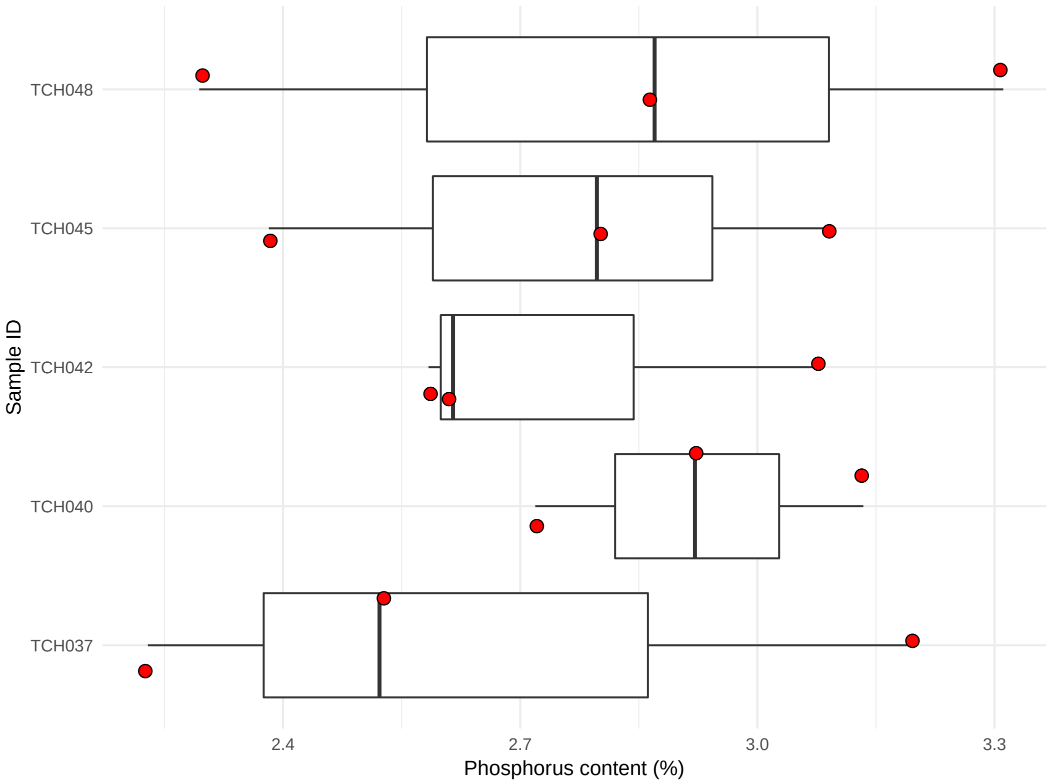

**Figure S4:** Variability in P content among three repeated samples taken from 5 random snowshoe hares after homogenization.

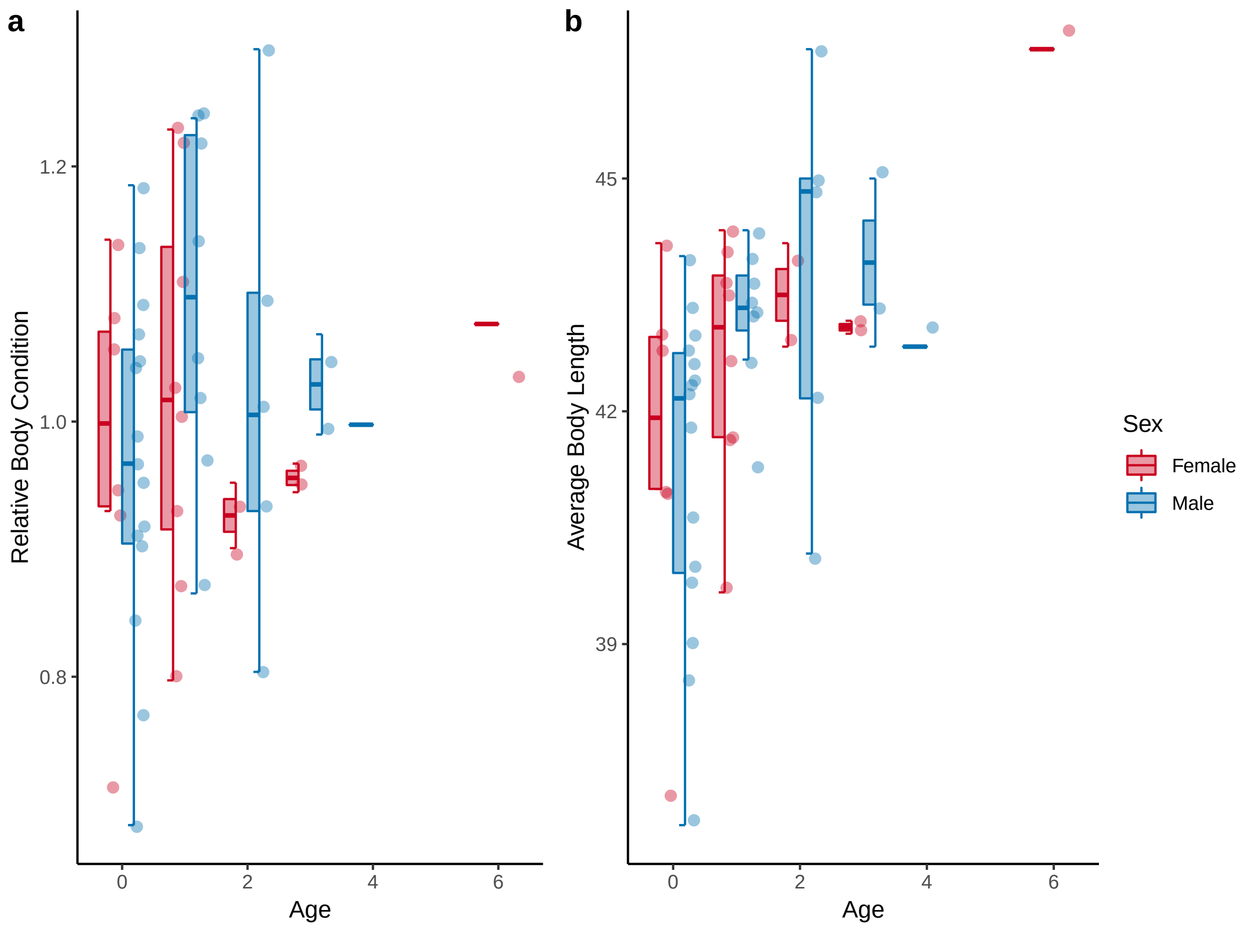

**Figure S5:** Variability in relative body condition (**a**) and average body length (**b**) with age, between the two sexes. Younger individuals appeared more variable than older ones. We found no evidence of a relationship between these variables through our statistical analyses.

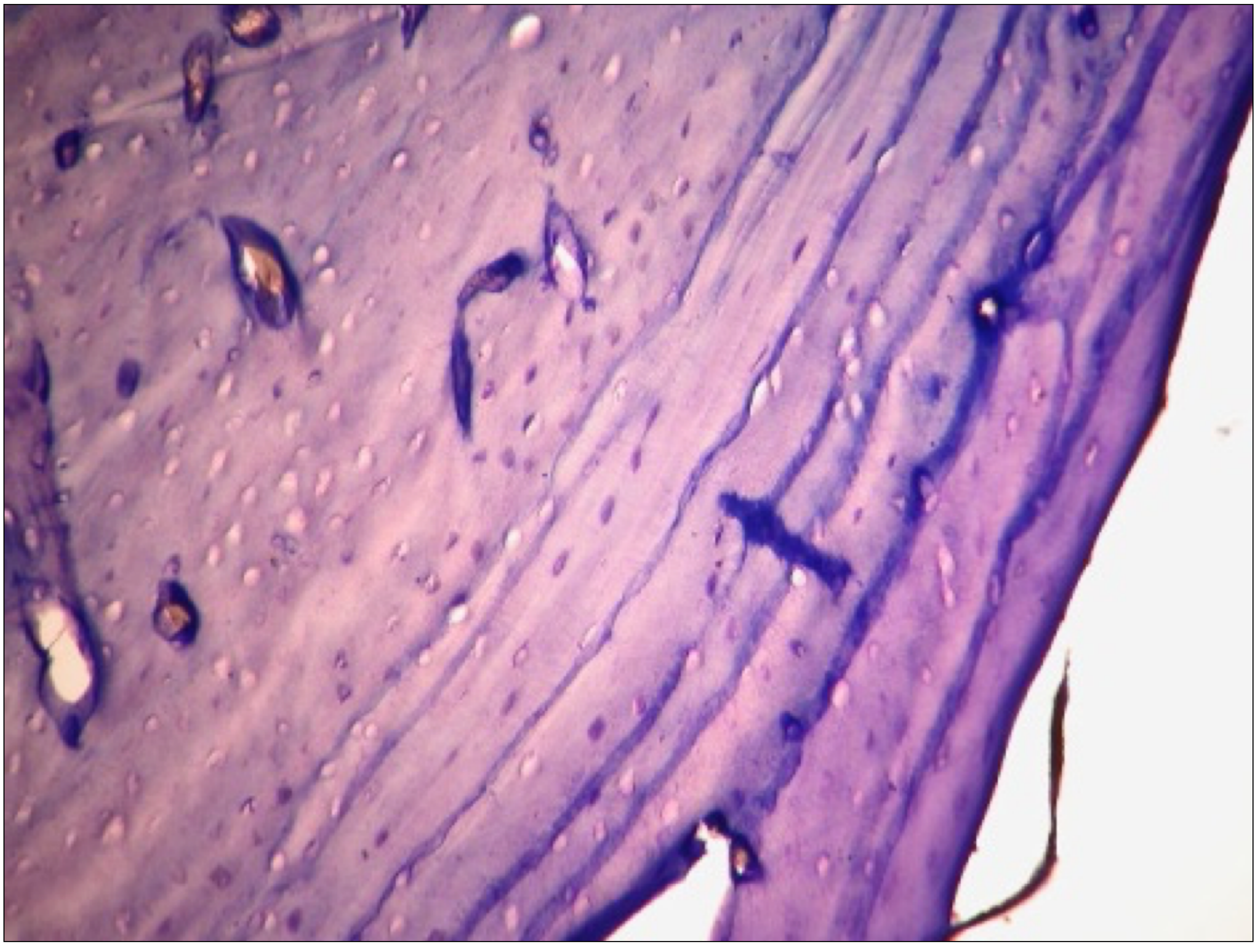

**Figure S6:** Histological preparation of mandibular section from hare TCH027, magnified 160X, showing the side of the mandible near the inferior surface. Estimated age for this individual is 6 years (i.e., 6 winters survived).

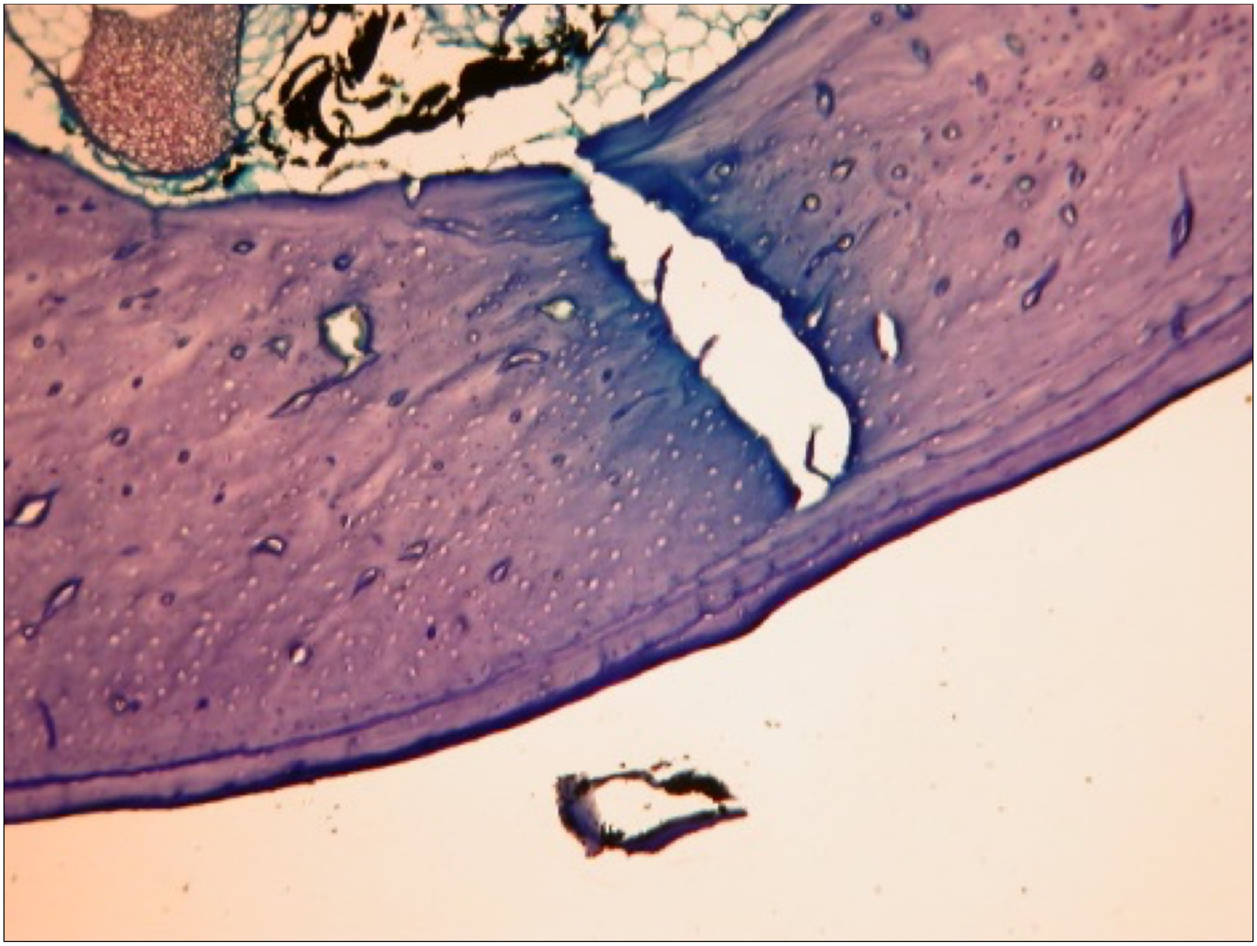

**Figure S7:** Histological preparation of mandibular section from hare TCH005, magnified 60X, showing the side of the mandible near the inferior mandible surface. In this case, age assessment is more difficult than in Figure S6. As the pattern is not so clear, a conservative age estimate would assign to this individual an age of 1 (i.e., 1 winter survived), with a range of 1–2 winters survived.
